# Not quite lost in translation: Mistranslation alters adaptive landscape topography and the dynamics of evolution

**DOI:** 10.1101/2022.12.15.520619

**Authors:** Michael Schmutzer, Andreas Wagner

**Affiliations:** Department of Evolutionary Biology and Environmental Studies, University of Zürich, Zürich, Switzerland; Swiss Institute of Bioinformatics, Lausanne, Switzerland; Santa Fe Institute, Santa Fe, New Mexico, USA

## Abstract

Mistranslation – the erroneous incorporation of amino acids into nascent proteins – is a source of protein variation that is orders of magnitude more frequent than DNA mutation. Like other sources of nongenetic variation, it can affect adaptive evolution. We study the evolutionary consequences of mistranslation with experimental data on mistranslation rates applied to three empirical adaptive landscapes. We find that mistranslation generally flattens adaptive landscapes by reducing the fitness of high fitness genotypes and increasing that of low fitness genotypes, but it does not affect all genotypes equally. Most importantly, it increases genetic variation available to selection by rendering many neutral DNA mutations non-neutral. Mistranslation also renders some beneficial mutations deleterious and vice versa. It increases the probability of fixation of 3 to 8 percent of beneficial mutations. Even though mistranslation increases the incidence of epistasis, it also allows populations evolving on a rugged landscape to evolve modestly higher fitness. Our observations show that mistranslation is an important source of non-genetic variation that can affect adaptive evolution on fitness landscapes in multiple ways.

## Introduction

Living organisms are no clockworks. The mechanisms by which they convert the information stored in their genes into phenotypes are imprecise and error-prone. As a result, even genetically identical individuals raised in the same environment vary in traits that affect fitness (Kotte et al. 2014; Solopova et al. 2014; Ackermann 2015; Govers et al. 2017; van Boxtel et al. 2017). The resulting nongenetic variation in fitness cannot be inherited with high fidelity (if at all). Any selective advantages or disadvantages it may cause will thus generally disappear within one generation (Bonduriansky and Day 2018).

Nongenetic fitness variation is ubiquitous and present at all levels of biological organisation, from the structure of macromolecules to the behaviour of whole organisms. Molecular examples of nongenetic fitness variation include gene expression noise (McAdams and Arkin 1999; Elowitz et al. 2002; Bódi et al. 2017), errors in transcription (Giacomelli et al. 2007; Traverse and Ochman 2016), and multiple alternative states in regulatory systems (Espinosa-Soto et al. 2011; van Heerden et al. 2014; Bruggeman and Teusink 2018) that underlie behaviours such as bet-hedging (Solopova et al. 2014; van Boxtel et al. 2017; Carey et al. 2018). Nongenetic fitness variation also exists in the phenotypes of multicellular organisms. Examples include random variation in behaviour, for example in the silk-spinning behaviour of fly larvae (Eberhard 1990), and the web-weaving of orb spiders (Eberhard 2000). Similar nongenetic variation is also documented in the nervous systems of mammals (Faisal et al. 2008). Here, we focus on mistranslation, i.e. the incorporation of amino acids different from those encoded in mRNA into a protein (Mordret et al. 2019), and its role for adaptive evolution.

Even though nongenetic fitness variation is not highly heritable, it can affect the rate of adaptation (Whitehead et al. 2008; Frank 2011; Rajon and Masel 2011; Draghi 2018; Rocabert et al. 2020). On the one hand, it can decrease the power of selection to act on genetic variation, for example by increasing the influence of genetic drift on a population (Wang and Zhang 2011; Mineta et al. 2015). On the other hand, nongenetic fitness variation can increase the fitness effects of some mutations. Specifically, it may increase the probability that beneficial mutations reach fixation, and it may increase the rate at which deleterious mutations are driven to extinction. In consequence, it may accelerate a populations’s rate of adaptation (Tănase-Nicola and ten Wolde 2008; Zhang et al. 2009; Frank 2011; Rajon and Masel 2011; Rocabert et al. 2020). For example, a yeast strain with higher variation than another strain in the expression of a protein conferring resistance to the antifungal drug fluconazole derives a larger fitness benefit from mutations in this protein, and thus evolves high resistance faster (Bódi et al. 2017). More generally, because of conflicting pertinent evidence, it is unclear whether nongenetic fitness variation is more likely to hinder or facilitate adaptation (Draghi 2018).

Whether nongenetic fitness variation benefits adaptation may depend on the dimensionality of a trait (Frank 2011; Rocabert et al. 2020). For example, gene expression noise may be able to accelerate adaptation in the expression level of a single gene (a single dimension; (Bódi et al. 2017)), but it might also slow down the evolution of a regulatory network with multiple genes (many dimensions). The reason is that any benefit derived from the expression level of one particular gene may be eliminated by detrimental expression noise in other genes. This ‘cost of complexity’ (Rocabert et al. 2020) may prevent nongenetic fitness variation from accelerating adaptation (Frank 2011; Rocabert et al. 2020).

A useful framework to investigate how high dimensional traits affect evolution is that of adaptive landscapes. Adaptive landscapes map genotypes within a collection or ‘space’ of genotypes onto fitness or some proxy thereof. They are also powerful conceptual tools for studying long-term evolution (Wright 1932; Wu et al. 2016; Aguirre et al. 2018; Ferretti et al. 2018; Bataillon et al. 2022). Because genotype spaces are high-dimensional discrete spaces, they have geometrical properties different from those of low-dimensional continuous spaces (Aguirre et al. 2018). Their high dimensionality itself has evolutionary consequences, for example when different mutations affect fitness non-additively (epistatically). Such epistasis can lead to rugged landscapes with many local peaks (Wu et al. 2016; Zagorski et al. 2016). In low dimensional spaces, epistasis can be an important barrier to adaptation, as populations can get stuck at local fitness optima. Higher dimensionality can create opportunities for escaping or bypassing local optima through ‘extra-dimensional bypasses’ in the higher dimensions (Wu et al. 2016; Zagorski et al. 2016). Thus a population evolving on a high dimensional, rugged landscape can reach higher fitness genotypes than on a low-dimensional landscape (Wu et al. 2016; Zagorski et al. 2016). Altogether, investigating how nongenetic fitness variation modifies adaptive landscapes may help address long-standing questions about how such variation can change adaptation.

In this study we focus on the effect of mistranslation on adaptive landscapes. Mis-translation is an unavoidable property of the translation machinery (Ribas de Pouplana et al. 2014), which erroneously incorporates amino acids into nascent proteins at rates between 10^−5^ to 10^−2^ per amino acid (Ribas de Pouplana et al. 2014; Mordret et al. 2019). As a result, about 10% of all proteins in *E. coli* will have an amino acid sequence that is different from their genetically encoded one (Ellis and Gallant 1982; Ruan et al. 2008). Not all amino acid substitutions are equally likely, and some codons have much higher mistranslation rates than others (Mordret et al. 2019).

For three reasons, mistranslation is highly useful to investigate the evolutionary impact of nongenetic fitness variation. First, the process linking a DNA genotype to a set of polypeptides produced through error-prone translation is well understood (Ribas de Pouplana et al. 2014; Mordret et al. 2019). The second reason stems from recent breakthroughs in high-throughput fitness assays (Blanco et al. 2019), which allow the mapping of large experimentally determined adaptive landscapes containing up to more than 10^5^ polypeptide sequences (Wu et al. 2016; Lite et al. 2020). Such landscapes permit modeling the fitness consequences of producing mistranslated polypeptides. Third, mistranslation can affect the course of adaptation. Even though it detrimentally impacts cell physiology and fitness (Javid et al. 2014; Ribas de Pouplana et al. 2014; Bratulic et al. 2017; Samhita et al. 2021), mistranslation can be beneficial for adaptation. For example, higher rates of mistranslation increase resistance to the antibiotic rifampicin in mycobacteria (Javid et al. 2014). Increased mistranslation of the *E. coli* antibiotic resistance gene TEM-1 *β*-lactamase facilitates the evolution of protein stability (Bratulic et al. 2015), and helps purge deleterious mutations (Bratulic et al. 2017; Zheng et al. 2021).

Here we build a quantitative model of the fitness effects of mistranslation that draws on experimental measured mistranslation rates (Mordret et al. 2019) and experimentally characterised high-dimensional adaptive landscapes (Wu et al. 2016; Lite et al. 2020). We use this model to investigate how mistranslation affects adaptive evolution on these landscapes. We quantify how frequently mistranslation facilitates or hinders the fixation of beneficial mutations, and find that mistranslation for the greater part hinders their fixation. Overall, mistranslation changes the paths available to populations evolving from low to high fitness genotypes, with many paths closing or becoming more difficult to traverse. The net effect of these changes is that populations can reach high fitness genotypes slightly better with mistranslation. We find that mistranslation improves the outcome of adaptation on two out of three adaptive landscapes we study. In addition, we find that mistranslation creates fitness differences between synonymous mutations, which can help improve adaptation among high fitness genotypes.

## Results

Mistranslation causes the fitness of a cell expressing a given genotype to depend not only on the protein encoded in the nucleotide sequence of the genotype (fig. 1A), but also on mistranslated variants of the same protein (fig. 1B). Because translation errors are random, mistranslation also causes variation between cells in the kind of protein variants they contain. These protein variants may have different activities, and thus cause variation in fitness among genetically identical cells.

**Fig. 1.**
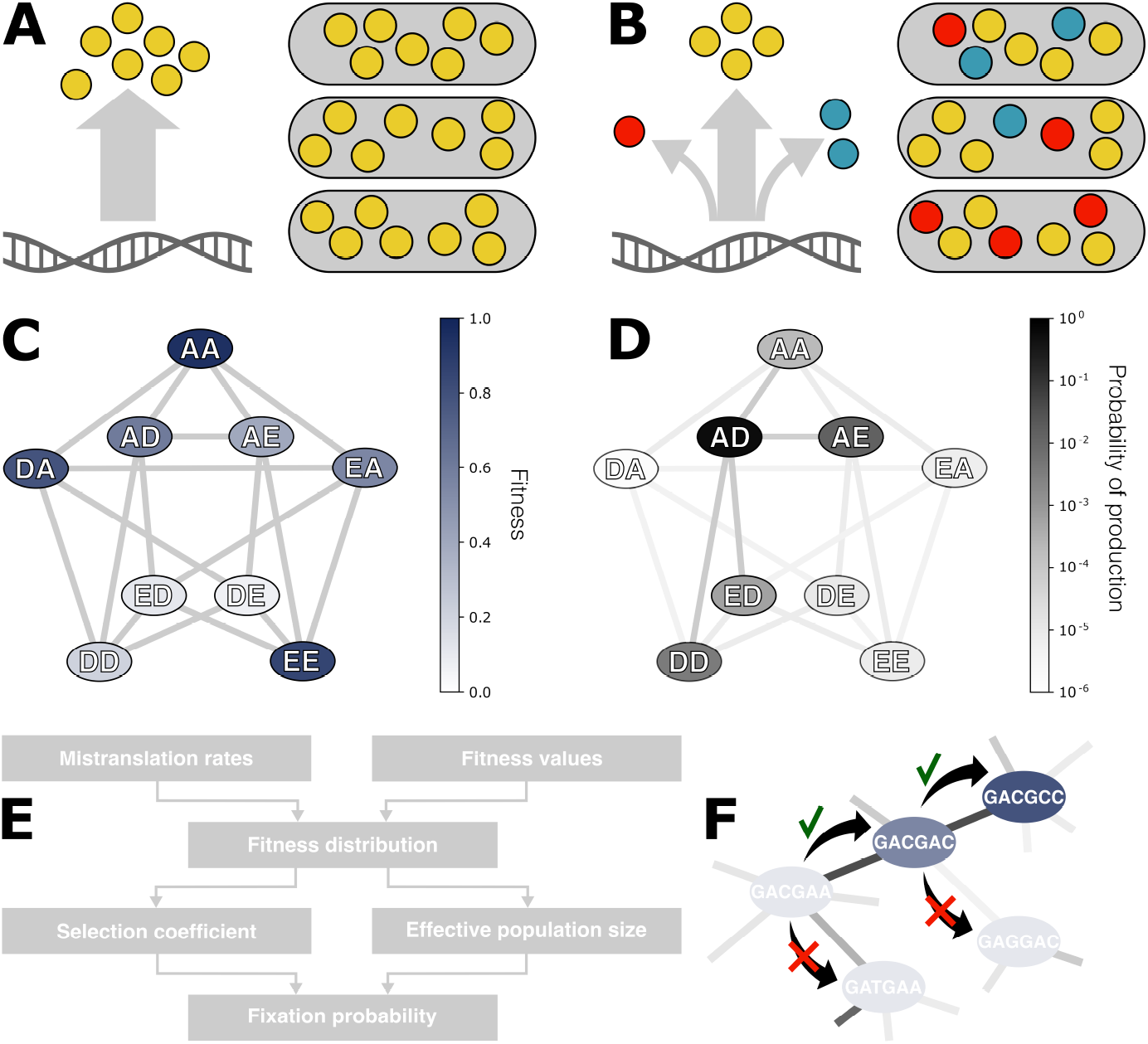
Modelling evolution under the influence of mistranslation. A) In the absence of mistranslation, cells produce proteins with exactly the amino acid sequence encoded in their genes. (B) Mistranslation causes the production of several protein variants (coloured circles) in different proportions (arrow thickness) from the same genotype. These protein variants do not necessarily have the same activity and can affect the fitness of a cell. Because mistranslation is a random process, cells with identical genotypes may still vary in protein variant composition, and thus potentially also in fitness. (C) We determine the effects of mistranslation from experimental data that quantified the fitness (or its proxy) of genotypes in an adaptive landscape at the polypeptide sequence level, as illustrated here with a toy example. Each polypeptide sequence (ellipses, linked to polypeptides that differ by one amino acid with grey edges) is encoded by a set of synonymous nucleotide sequences. (D) We simulate mistranslation using experimentally measured mistranslation rates (Mordret et al. 2019). Each codon differs in its mistranslation rate, and thus each nucleotide sequence has a different propensity to produce alternative protein variants. For example, a nucleotide sequence encoding the polypeptide ‘AD’ (black ellipse) has a higher probability of producing certain polypeptide sequences with a single amino acid substitution than others (e.g. ‘AE’ is more likely than ‘ED’ in this example), and is even less likely to produce a protein variant with a double substitution (‘DA’). We calculate the composition of protein variants expected under mistranslation for a given nucleotide sequence, and then use the fitness measurements of these protein variants from (C) to estimate a cell’s expected fitness. (E) Model work flow. Using the experimental measurements of fitness and mistranslation rates, we estimate the fitness distributions of a cell expressing a given genotype and of a cell expressing a mutant genotype. From these two fitness distributions we calculate the mutant’s selection coefficient and the effective population size, which is influenced by mistranslation (Materials and Methods). We then use these estimates and Kimura’s (Kimura 1962) equation to estimate the fixation probability of the mutation. (F) We simulate evolution at the nucleotide sequence level in a weak mutation-strong selection regime, where new mutant genotypes fix rapidly compared to the time they need to originate. In this regime populations harbour only one allele at the loci we simulate for most of the time. Consequently, evolution resembles a ‘walk’ from one genotype (ellipses, coloured by fitness, edges connect genotypes differing in one nucleotide position) to the next. Each step in the walk is equivalent to the fixation of a novel genotype. In our model, mutations change a single nucleotide position (black arrows). Every mutation is equally likely to occur, but its fixation probability (edge shading) depends on its fitness relative to the current genotype. Mutations that decrease fitness are likely (but not necessarily) lost (red crosses), while mutations that increase fitness are likely to reach fixation (green check-marks).

To model these effects of mistranslation on fitness, we draw on experimentally measured mistranslation rates (Mordret et al. 2019), and three different empirical adaptive landscapes (Wu et al. 2016; Lite et al. 2020). Details on these data and data curation are available in supplementary section 3. In short, the mistranslation rates are estimated from the frequencies of different protein amino-acid sequence variants in *E. coli* detected through mass-spectrometry (Mordret et al. 2019). The first of the landscapes comprises measurements of the binding affinity of almost 160,000 genotypic variants of the streptococcal immunoglobulin-binding protein (GB1), which can bind antibodies (Wu et al. 2016). In using data from this landscape, we follow other authors in considering strong molecular binding as a proxy for high fitness (Natarajan et al. 2013; Olson et al. 2014; Sarkisyan et al. 2016; Aguilar-Rodríguez et al. 2017). We will refer to this landscape as the antibody-binding landscape. The second and third landscapes comprise data on the growth rate of *E. coli* cells transformed with almost 8,000 variants of a toxin-antitoxin system (Lite et al. 2020). Specifically, these are variants of the ParD3 antitoxin, part of a toxin-antitoxin system that targets topoisomerases (Jiang et al. 2002; Yuan et al. 2010). The ParD3 antitoxin variants were either expressed together with its native ParE3 toxin, or with its close homologue ParE2, resulting in two fitness measurements per antitoxin variant. We refer to these landscapes as the toxin-antitoxin landscapes, and differentiate them where necessary by the target toxins E3 or E2.

We combine data from these experimentally mapped adaptive landscapes (fig. 1C) with experimentally measured mistranslation rates (fig. 1D) to quantify the average change in fitness and the variation in fitness between individuals that is caused by mistranslation. The same data also allows us to quantify how mistranslation changes the outcome of adaptive evolution. To this end we simulate adaptive evolution on the adaptive landscapes defined by the data. Adaptive evolution occurs at the nucleotide sequence level, and can be influenced by the mistranslation of individual codons on the mRNA into different amino acids. Our model allows us to estimate the protein variant compositions of a population of cells expressing a given nucleotide sequence (fig. 1D) from these codon mistranslation rates (Mordret et al. 2019). We then estimate the fitness distribution of cells in the population based on the fitness of these protein variants (fig. 1D). Using the fitness distributions (fig. 1E) of a population carrying a given (‘wild-type’) genotype and another carrying a mutant genotype, we can then estimate the selection coefficient of the mutant, which we define as the relative difference in mean fitness between these genotypes. We also estimate the effective population size, which can be affected by the mistranslation rate (Wang and Zhang 2011). These two quantities determine the fixation probability of the mutant genotype when introduced at a low frequency into a population consisting only of wild-type genotypes.

Because the experimentally determined landscapes we consider (Wu et al. 2016; Lite et al. 2020) are based on a small number of nucleotide sites, we assume that adaptive evoluiton occurs in the weak mutation, strong selection regime (Gillespie 1983; Gillespie 1984). In this regime, only one mutation segregates at any point in time, and adaptive evolution effectively becomes an adaptive walk in which a population steps from one genotype to the next (Wright 1932; Maynard Smith 1970; Gillespie 1984) (fig. 1F). The probability of each step is defined by the fixation probabilities we calculated from the data. While simulating such adaptive walks, we quantify evolutionary time through the number of mutations that occurred since the beginning of the walk. We keep track of the small fraction of these mutations that go to fixation.

Using such adaptive walks, we study how mistranslation affects adaptive evolution on three adaptive landscapes that are combinatorially complete for all 20 amino acids (Materials and Methods, (Wu et al. 2016)).

### Mistranslation generally flattens adaptive landscapes

We first investigate the impact of mistranslation on properties of genotypes that are important for adaptive evolution. Specifically, we assess how mistranslation affects (i) the expected (mean) fitness of genotypes, (ii) the effective population size (Materials and Methods), (iii) the fitness effects of mutations, and (iv) the fixation probabilities of mutations. We characterise the effect of mistranslation for 10^4^ randomly chosen genotypes from each landscape and at three population sizes *N* = 10^4^, 10^6^ and 10^8^. We find that many, but not all, of the effects of mistranslation are subtle, because the mistranslation rates we use are realistically small (Mordret et al. 2019). Our results in this section are very similar for all three landscapes, therefore we will focus on the antibody-binding landscape (see supplementary fig. S1 and S2 for the toxin-antitoxin landscapes).

The production of mistranslated polypeptides can change the mean fitness of a population of cells, because these mistranslated polypeptides vary in their effect on fitness. Mistranslation increases the fitness of some genotypes, and decreases the fitness of other genotypes (fig. 2A). The direction of this change in fitness depends on a genotype’s fitness in the absence of mistranslation. Mistranslation tends to increase the fitness of genotypes with a fitness below the mean fitness of 0.08 in the antibody-binding landscape. In contrast, mistranslation tends to decrease the fitness of genotypes with higher fitness. In other words, mistranslation causes the landscape to become ‘flattened’, with high fitness genotypes decreasing in fitness due to mistranslation, and low fitness genotypes increasing in fitness. As a result of this flattening, the slope of the correlation *m* between the fitness without and with mistranslation is less than one (*m* = 0.988, Pearson’s *r* = 1.00, *p* ≪ 10^−300^, difference between observed slope and a slope of one *t* = 266, *p* ≪ 10^−300^).

**Fig. 2.**
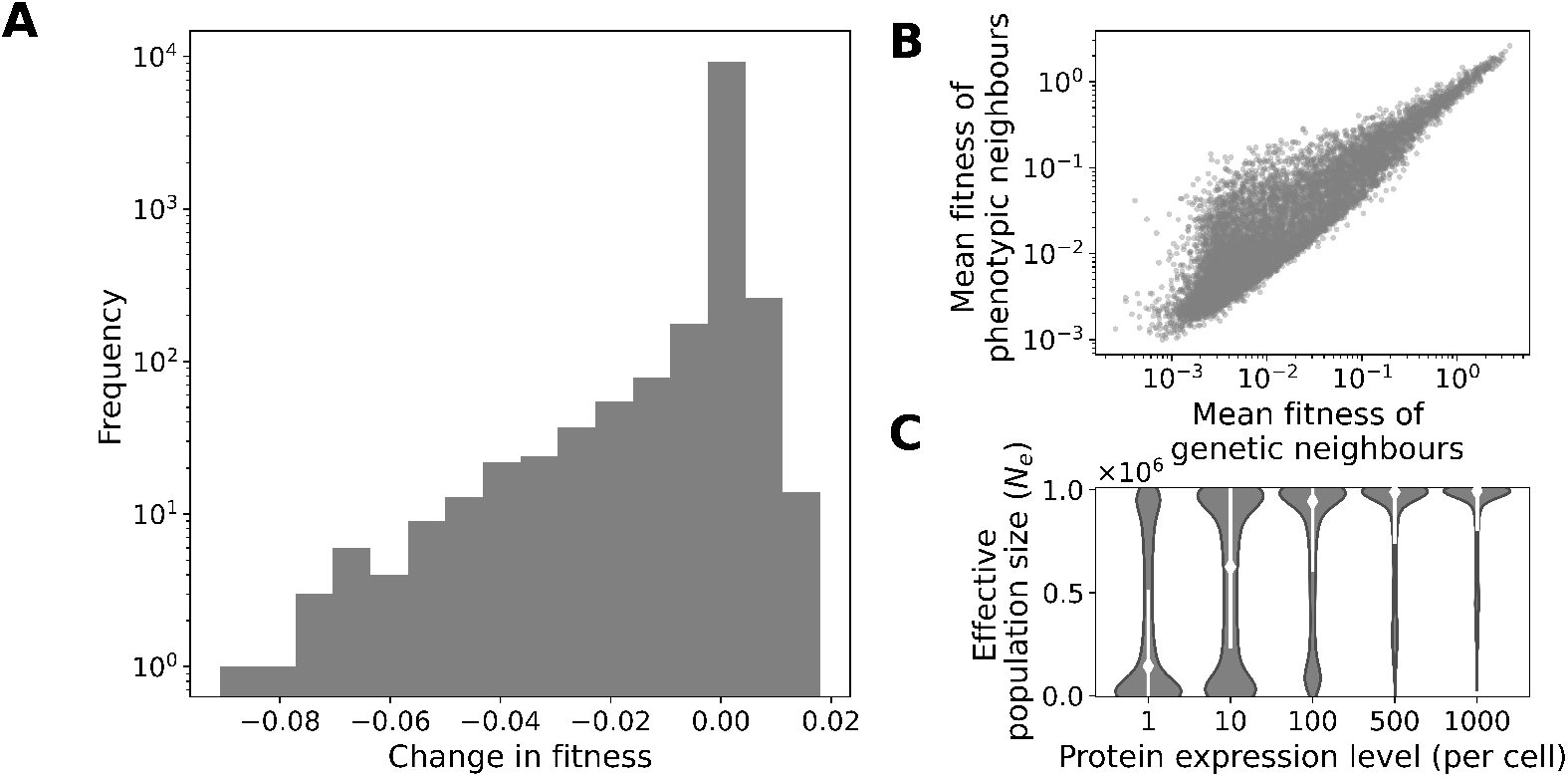
Mistranslation flattens the fitness distribution of the antibody-binding landscape. A) Distribution of the changes in fitness due to mistranslation of 10^4^ randomly chosen genotypes. B) The mean fitness of a genotype’s phenotypic neighbours (polypeptide sequences that differ in one amino acid position from the genotype, vertical axis) and the mean fitness of its genotypic neighbours (genotypes with a single nucleotide change, horizontal axis) are positively correlated. C) Mistranslation reduces the effective population size. White diamonds and lines show the median effective population size and the standard deviation, respectively. The violin plots show a Gaussian kernel density estimate of the distribution of effective population sizes. In all panels results are shown for the same set of 10^4^ randomly chosen genotypes from the antibody-binding landscape, at a population size of *N* = 10^6^, and an expression level of one protein per cell (unless where stated otherwise).

We also find that the fitness values of the phenotypic neighbours of a given genotype (the polypeptide sequences that differ by one amino acid from the polypeptide encoded by the genotype) are correlated with the fitness values of its genotypic neighbours (fig. 2B, Kendall’s *τ* = 0.72, *p* ≪ 10^−300^, *n* = 10^4^). In order words, the nongenetic fitness variation created through mistranslation is related to the genetic fitness variation present in the immediate neighbourhood of a genotype in sequence space. This is a necessary condition for mistranslation to facilitate adaptative evolution (Ancel and Fontana 2000; Frank 2011; Rocabert et al. 2020).

### Mistranslation strengthens genetic drift

Mistranslation causes variation in protein composition between genetically identical cells, which can induce small variation in fitness between cells. The degree of fitness variation between individuals depends on a protein’s expression level. When few copies of a protein exist in a cell, a single mistranslated copy can affect fitness more substantially than when many copies exist. Consequently, fitness will vary more strongly between cells that produce fewer copies of a protein. This kind of nongenetic fitness variation can weaken selection, an effect that can be quantified as a decrease in the effective population size (Materials and Methods, (Wang and Zhang 2011)). To determine the strength of this effect, we thus quantify the effect of protein expression level on the effective population size, and do so for protein expression levels between 10^0^ to 10^3^ proteins per cell.

As predicted, we find that mistranslation causes a greater reduction in the mean effective population size at lower than at higher expression levels (fig. 2C). For example, mistranslation causes effective population size in a population of *N* = 10^6^ individuals to decrease on average by 66 percent to *N_e_* = 3.4 *×* 10^5^ (*±*3.7 *×* 10^5^ standard deviation) for a protein expressed at one copy per cell, but by only 11 percent to *N_e_* = 8.9*×*10^5^ (*±*2.0*×*10^5^) for a protein expressed at 10^3^ copies. Effective population size also varies greatly even within one expression level depending on the protein genotype that is considered. The reason is that different genotypes vary in their rates of mistranslation and the fitness values of their phenotypic neighbours. Similar observations hold for the toxin-antitoxin landscapes (supplementary fig. S1 and S2). Throughout the rest of this paper we will distinguish between the population size *N* and the effective population size *N_e_ ≤ N* that is a consequence of mistranslation.

### Mistranslation reduces neutrality and can cause deleterious mutations to turn beneficial and vice versa

Changes in the fitness of genotypes may also affect the fitness of mutations. To determine how strongly mistranslation affects the fitness of mutations, we sample 10^4^ genotypes at random from the antibody-binding landscape, and chose one of their mutational neighbours, i.e. a genotype that differs from the focal genotype by one nucleotide substitution. We classify the fitness effects of these mutants as beneficial, deleterious or (nearly) neutral (if 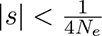 (Charlesworth 2009)), compared to the wild-type genotype, both in the presence and absence of mistranslation (fig. 3A).

**Fig. 3.**
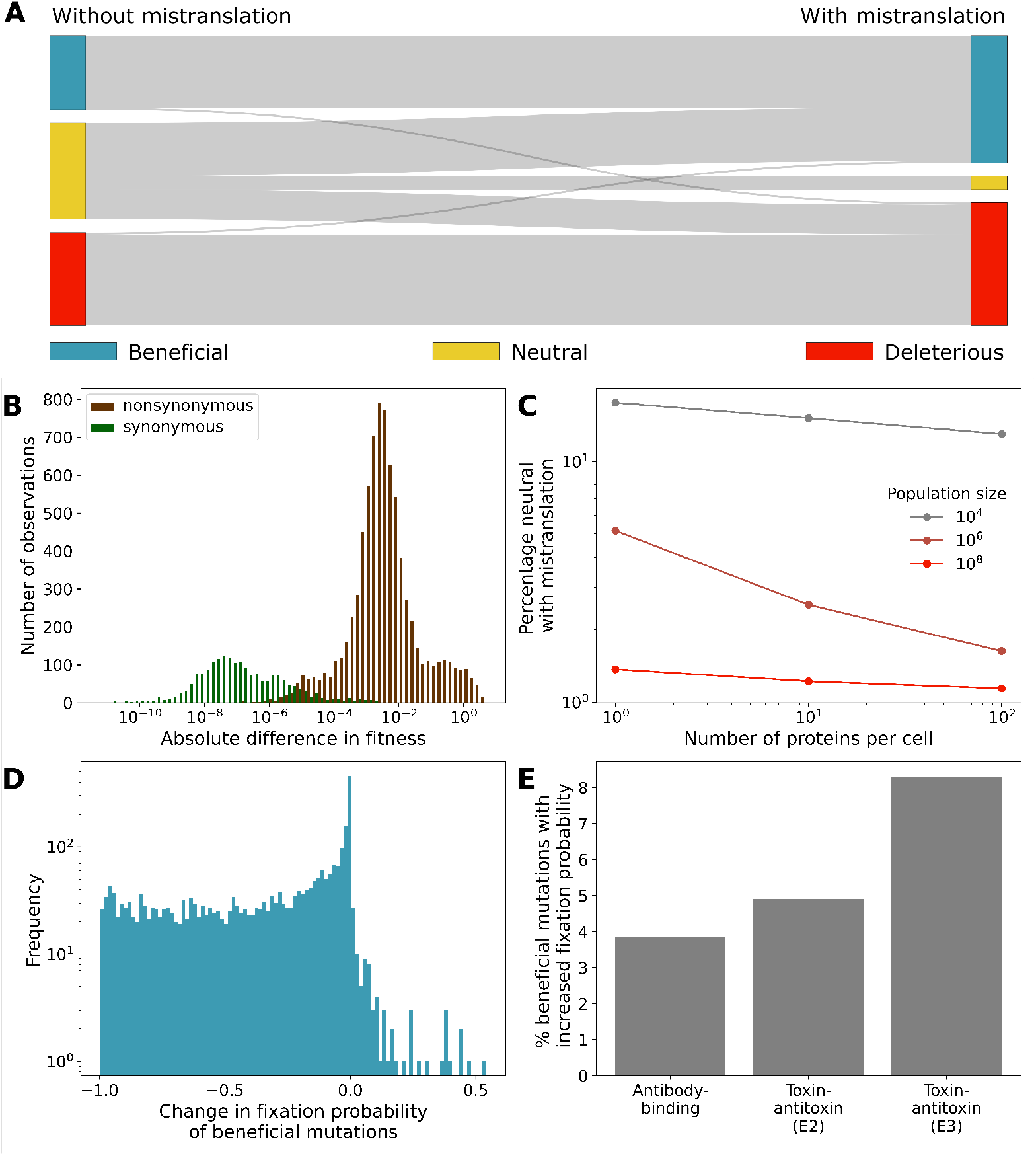
Mistranslation changes the fitness effects of some mutations. A) Changes due to mistranslation in the proportions of mutations classified according to their fitness effects as beneficial, (nearly) neutral, or deleterious. B) Distributions of the absolute differences in mean fitness between both synonymous (green) and nonsynonymous (brown) pairs of neighbouring genotypes sampled from the antibody-binding landscape in the presence of mistranslation at an expression level of one protein per cell. Only nonzero fitness differences are shown. C) Percentage of mutations classified as neutral in the presence of mistranslation for multiple combinations of the population size *N* and the protein expression level, both of which impact the effective population size *Ne* (and thus neutrality). D) Distribution of the change in fixation probabilities of beneficial mutations due to mistranslation. Only those mutations are shown that are beneficial both with and without mistranslation. The horizontal axis shows the difference in fixation probability due to mistranslation, with zero denoting no change. For example, changes in fixation probability close to negative one signify mutations that are almost certain to fix without mistranslation (*u_fix_ ≈* 1), but have almost no chance of reaching fixation with mistranslation (*u_fix_ ≈* 0). E) Percentage of beneficial mutations that increase in fixation probability due to mistranslation for each landscape. In all panels (except where stated otherwise), results are shown for the same set of 10^4^ randomly chosen genotypes from the antibody-binding landscape as in fig. 2, and randomly chosen one-step mutational neighbours, at a population size of *N* = 10^6^, and an expression level of one protein per cell. The same parameters apply for the two toxin-antitoxin landscapes in (E).

This analysis shows that mistranslation can cause substantial changes in fitness effects. In particular, mistranslation dramatically reduces the number of nearly neutral mutations. For example, at a population size of *N* = 10^6^ and a low expression level of one copy per cell, 37% of mutations in the antibody-binding landscape are nearly neutral in the absence of mistranslation, but only 5% are nearly neutral in the presence of mistranslation and low expression. 56% of the mutations that are nearly neutral in the absence of mistranslation are synonymous mutations, and 74% of these synonymous nearly neutral mutations come under selection in the presence of mistranslation. 20% of nonsynonymous mutations are neutral in the absence of mistranslation, and all of these mutations become non-neutral under mistranslation. Moreover, the fitness differences between nonsynonymous mutations are orders of magnitude larger than the differences between synonymous mutations induced by mistranslation (fig. 3B). This loss of nearly neutral mutations is influenced both by the population size and the protein expression level (fig. 3C). The loss of nearly neutral mutations also severely affects the size of nearly neutral networks (supplementary section 4).

In addition, we observe that 2.5% of mutations that are beneficial in the absence of mistranslation become deleterious in its presence, and 2.1% of mutations change from deleterious to beneficial (fig. 3A). The reason is that the flattening of the fitness landscape caused by mistranslation does not happen to the same extent for all genotypes, because genotypes differ both in their rate of mistranslation and their genetic neighbourhoods. For example, a mutation that is beneficial in the absence of mistranslation may become deleterious because it has a higher mistranslation rate than the wild-type, or because the mutation increases the probability of producing a highly deleterious protein variant through mistranslation.

Even when mistranslation does not change the fitness effect of a mutation from beneficial to deleterious, it can affect the mutation’s fixation probability (fig. 3D). For example, in the antibody-binding landscape 95.7 percent of beneficial mutations experience a reduction in this probability, and only 3.9 percent of mutants experience an increase. In this respect, the toxin-antitoxin landscapes are different from the antibody-binding landscape. For example, the fraction of beneficial mutations whose fixation probability increases is much greater (8.3 percent) in the toxin-antitoxin (E3) landscapes (fig. 3E). These changes in the fixation probability of beneficial mutations are largely driven by reductions in the effective population size (supplementary fig. S3). For the deleterious mutations we examined, the fixation probability is largely unaffected with a mean increase of 2.81 *×* 10^−79^ *±* 1.65 *×* 10^−77^ in the antibody-binding landscape, and a mean increase of 6.69 *×* 10^−15^ *±* 3.64 *×* 10^−13^ in the E3 toxin-antitoxin landscape, but a mean decrease of 6.89 *×* 10^−101^ *±* 3.62 *×* 10^−99^ in the E2 toxin-antitoxin landscape (*N* = 10^6^).

### Mistranslation increases the ruggedness of landscapes

Given that mistranslation causes a flattening of adaptive landscapes, mistranslation may also change the ruggedness of a fitness landscape with multiple peaks, and thus also how accessible high fitness genotypes are. The reason is that in a rugged landscape, evolving populations are more likely to become stuck at peaks of low fitness (Poelwijk et al. 2007). Because mistranslation may ‘smooth over’ some of the roughness of a landscape, it may render high fitness genotypes genotypes more accessible through fitness increasing paths (Frank 2011). Intriguingly, the following analysis does not support this prediction. We estimate the ruggedness of the three landscapes with two complementary measures, the number of fitness peaks in a landscape and the frequency of different kinds of epistasis. For the purpose of our analysis, a fitness peak can consist of a single genotype or a network of nearly neutral genotypes that are single-nucleotide mutational neighbours of one another (supplementary section 3). We consider any such nearly neutral network a fitness peak if all single-step mutational neighbours of the genotypes in the network have a lower fitness.

Because the size of nearly neutral networks is affected by population size, we evaluate the number of fitness peaks at different population sizes. Mistranslation increases the number of fitness peaks (fig. 4A), except at low population sizes (supplementary fig. S4). Most of these additional fitness peaks arise because synonymous mutations come under selection in large populations.

**Fig. 4.**
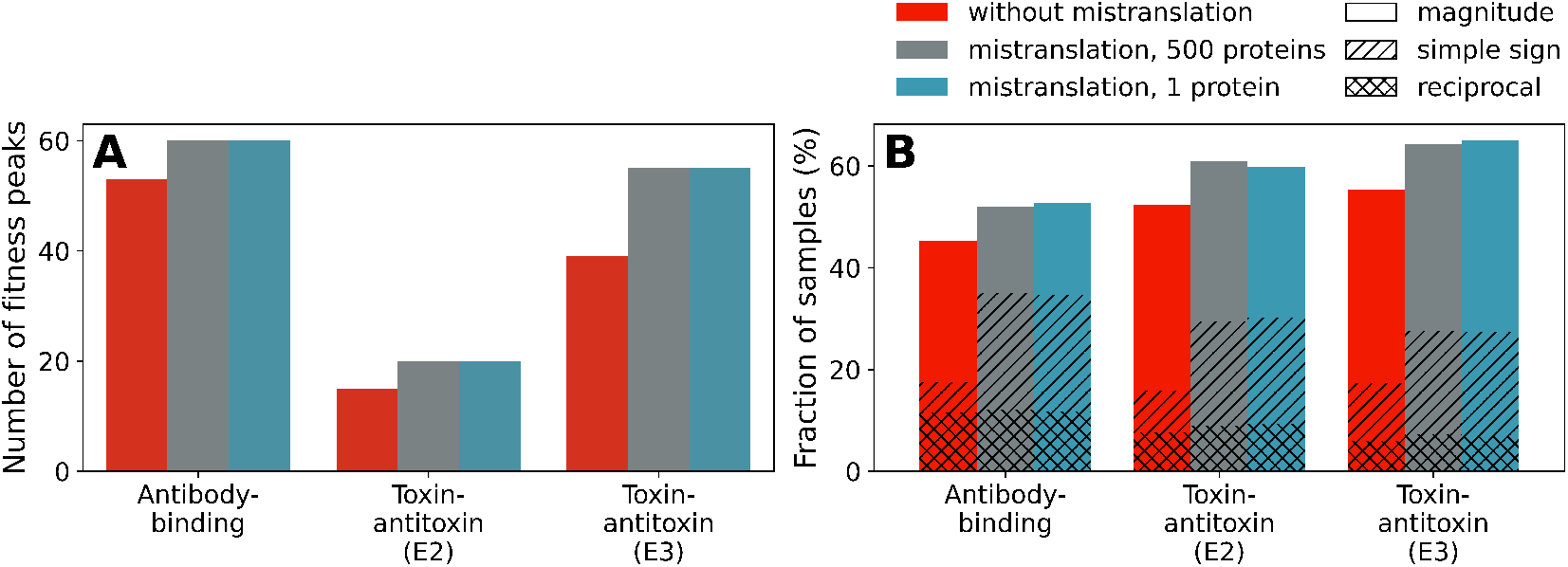
Mistranslation increases the ruggedness of all three landscapes. A) Number of fitness peaks among 10^5^ genotypes with the highest fitness in all three landscapes, without mistranslation (red), and with mistranslation at both high and low protein expression levels (500 proteins or one protein per cell, grey and blue, respectively). Fitness peaks are evaluated at large population sizes (*N* = 10^8^). B) Percentage of 10^4^ genotype samples classified as one of three kinds of epistasis. Each sample consists of a ‘square’ of four genotypes, two of which are double nucleotide mutants of each other, and the other two are single mutant intermediates. If the fitness effects of the double mutant is not additive with respect to the fitness effects of the single mutants, the two mutations are epistatic, with the kind of epistasis depending on the nature of the deviation from additivity (supplementary section 3, (Poelwijk et al. 2007), see also supplementary fig. S6). For all three landscapes, we determine epistasis for the same set of 10^4^ sequences without and with mistranslation, and in the latter case for high and low protein expression.

Another measure of ruggedness is the frequency of multiple kinds of non-additive (epistatic) interactions between pairs of single nucleotide mutants (Poelwijk et al. 2007). One can distinguish three kinds of such non-additivity (epistasis): Magnitude epistasis, simple sign epistasis and reciprocal sign epistasis. The latter two are associated with the existence of fitness valleys that may decrease the number of accessible (monotonically fitness-increasing) paths towards high fitness peaks (Poelwijk et al. 2007). Our analysis shows that mistranslation makes all kinds of epistasis more common, particularly magnitude and simple sign epistasis, and to a lesser extent also reciprocal sign epistasis (fig. 4B). These observations show that mistranslation increases the ruggedness of a landscape. Thus, if mistranslation increases the accessibility of high fitness peaks, it does not do so by landscape smoothing.

Taken together, the results presented in the current and the previous section suggest that mistranslation is not simply smoothing a landscape, but rather changing which paths from low fitness genotypes to high fitness genotypes are available to populations evolving on these landscapes. Some paths are closed off by mistranslation, for example because mistranslation has turned a beneficial mutation into a deleterious one. Alternatively, many paths become more difficult to traverse for evolving populations, for example because the fixation of the beneficial mutations that constitute these paths has become less probable. However, mistranslation also opens up new paths that arise due to neutral and deleterious mutations that become beneficial, and some rare paths become easier to traverse because a small fraction of beneficial mutations increase in fixation probability. Depending on where these novel paths lead in the adaptive landscape, mistranslation may affect the fitness attained by populations evolving on the landscape.

### Mistranslation helps evolving populations attain higher fitness on two landscapes

We next investigate how mistranslation influences adaptive evolution on all three adaptive landscapes. To this end, we choose 10^4^ starting genotypes from each landscapes at random among the viable genotypes that rank in the bottom 10% of the fitness distribution. We perform adaptive walks (Materials and Methods) for three population sizes (10^4^, 10^6^, 10^8^), and two levels of protein expression. Specifically, we consider expression levels of one and 500 proteins per cell, where 500 proteins is the median expression level in *E. coli* (Ishihama et al. 2008). We will refer to these expression levels as low and high expression respectively. We will refer to the number of mutations (both fixed and extinct) since the beginning of a walk as a proxy for evolutionary time.

We find that mistranslation facilitates adaptation in the antibody-binding landscape, but much less so or not at all in the two toxin-antitoxin landscapes. Specifically, at the end of adaptive walks on the antibody-binding landscape, populations with mistranslation attain slightly higher mean fitness, and this advantage increases with larger population sizes. For example, for populations with *N* = 10^4^ individuals and low expression, the average adaptive walk ends at genotypes that have 52.1 (*±* 26.8, one standard deviation) and 50.2 (*±* 27.7) percent of the maximum fitness, respectively, in the presence and absence of mistranslation. These differences are small but statistically significant (Dunn’s test with Bonferroni correction, *p ≤* 2.82 *×* 10^−9^).

At a population size of 10^8^ adaptive walks with mistranslation and low expression reach a fitness that is 2.9 percent higher than in its absence (50.2 *±* 26.1 and 47.3 *±* 27.2, percent of the maximum, respectively, with significant differences only between walks with and without mistranslation (Dunn’s test with Bonferroni correction, *p* = 4.70 *×* 10^−17^ or smaller). Compared to the antibody-binding landscape the effect of mistranslation on the outcome of adaptive walks on the toxin-antitoxin (E3) landscape is an order of magnitude smaller, and we do not observe a consistent pattern in the toxin-antitoxin (E2) landscape (supplementary table S1 and section 5).

Protein expression level has little effect on the outcome of adaptive evolution with mistranslation. For example, for adaptive walks on the antibody-binding landscape, populations with *N* = 10^8^ individuals reach 50.2*±*26.1 and 49.9*±*26.5 percent of the maximum fitness, a difference in outcome that is not statistically significant (Dunn’s test with Bonferroni correction, *p* = 1.00). Similar observations hold for other population sizes on the antibody-binding landscape and on the toxin-antitoxin landscapes (supplementary tables S1 and S2). However, protein expression level does affect how fast adaptive walks attain high fitness (supplementary section 6).

In addition, escapes from local fitness peaks play a relatively small role in the outcome of adaptive walks. In 23.0 percent of adaptive walks on the antibody-binding landscape, populations with low expression and small population sizes (*N* = 10^4^) escape from local fitness peaks through the fixation of slightly deleterious mutations. However, at the highest population size (*N* = 10^8^), we do not observe any such escapes from local fitness peaks in adaptive walks on any of the three adaptive landscapes. Consequently, escapes from local fitness peaks are not sufficient to explain the improved outcome of adaptive walks with mistranslation.

### Synonymous mutations facilitate continued adaptation at high fitness

Most of the fitness gains during adaptive walks happen within the first 10^3^ mutations of an adaptive walk. This increase is largely driven by nonsynonymous mutations. For example, nonsynonymous mutations contribute on average 99.98 *±* 0.03 percent of the change in fitness during walks with mistranslation on the antibody-binding landscape (*N* = 10^8^, high protein expression). However, evolution involving both selection and drift may continue for a long time among high fitness genotypes. Specifically, we hypothesise that mistranslation can lengthen adaptive walks. The reason is that high fitness genotypes subject to mistranslation largely produce deleterious protein variants, and selection may prefer codons associated with low mistranslation. Consequently, adaptive walks with mistranslation will tend to fix additional synonymous mutations that do not become fixed in walks without mistranslation.

To test this hypothesis, we study our adaptive walks in greater detail. In support of it, we find that in all three landscapes, adaptive walks with mistranslation fix significantly more mutations than walks without mistranslation, except at small population sizes and in walks on the antibody-binding landscape with mistranslation and low expression (supplementary table S3). The difference is small, partly because only few of a walk’s 10^5^ mutations go to fixation (mean number of fixations: 8.4*±*3.1 vs 7.7*±*3.1, fig. 5A, antibody-binding landscape, *N* = 10^8^, high expression, Wilcoxon test *z* = 16301820, *p* = 4.86 *×* 10^−57^, *n* = 10^4^). These differences come from an increase in the number of synonymous mutations going to fixation in adaptive walks with mistranslation. For example, on the antibody-binding landscape, without mistranslation and a population size of *N* = 10^8^, we observe on average only 3.0 *×* 10^−4^ (*±*1.7 *×* 10^−2^) synonymous and 7.7 (*±* 3.1) nonsynonymous mutations per adaptive walk. In contrast, with mistranslation at high expression we observe on average 0.6 (*±* 0.7) synonymous and 7.8 (*±* 3.0) nonsynonymous mutations per walk. In other words, mistranslation helps increase the number of synonymous substitutions by three orders of magnitude.

**Fig. 5.**
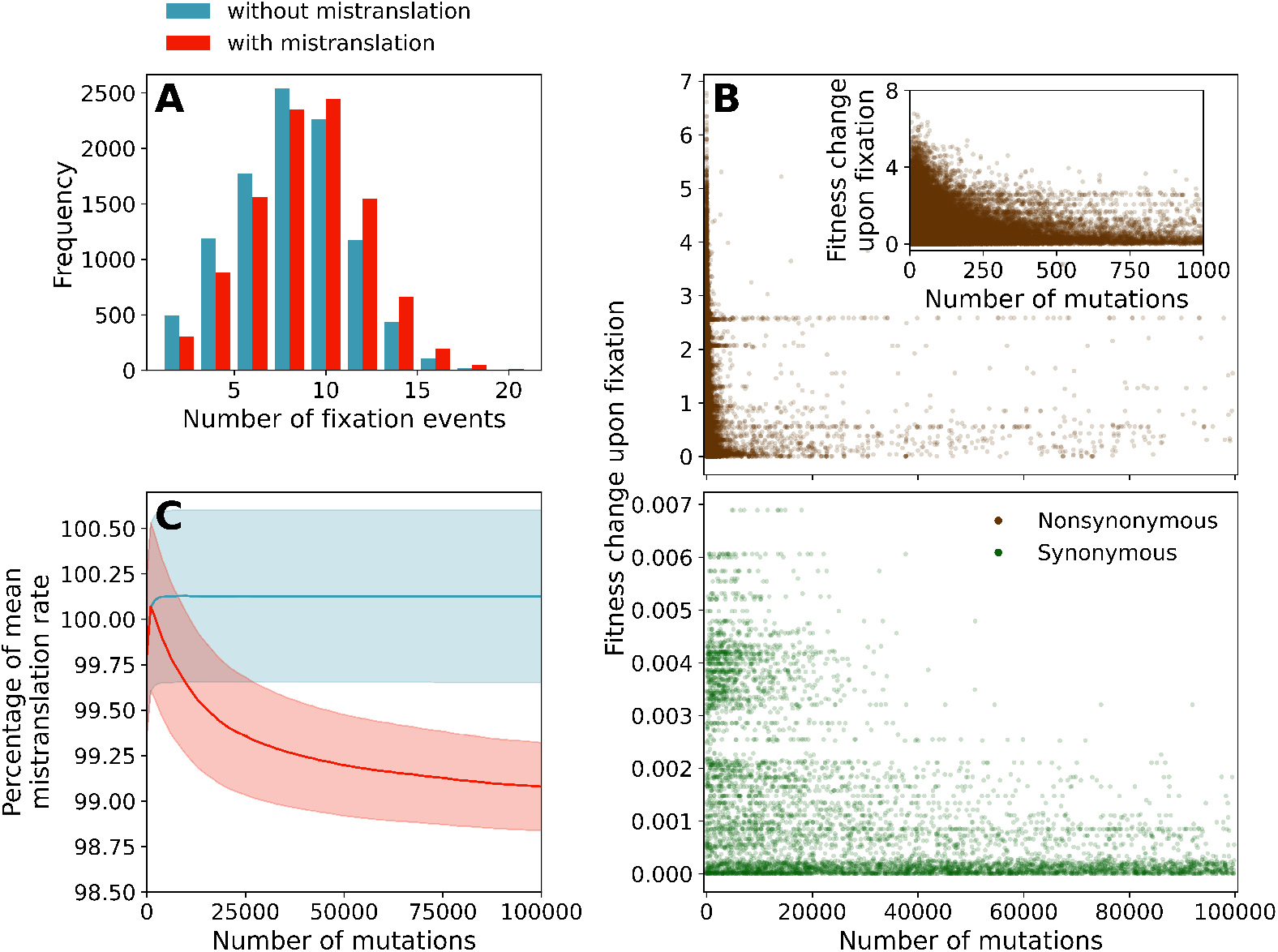
Mistranslation causes longer adaptive walks due to selection against high mistranslation rates acting on synonymous mutations. A) Distributions of the number of mutations that reach fixation during adaptive walks with (red) and without mistranslation (blue). B) Fitness effects (vertical axis) of all nonsynonymous (brown, upper panel) and synonymous (green, lower panel) mutations fixed during adaptive walks with mistranslation. The horizontal axis shows the number of mutations (both extinct and fixed) since the beginning of the adaptive walk. Inset: Fitness effects of nonsynonymous mutations that reach fixation within the first 1000 mutations of the adaptive walk. C) Mean and standard error (lines and shaded areas, respectively) of the mistranslation rates (red) and post hoc mistranslation rates (blue) of the genotypes encountered during adaptive walks with and without mistranslation rates, respectively. (Post hoc) mistranslation rates are shown as a percentage of the mean mistranslation rate of all genotypes in the antibody-binding landscape (100 percent is equivalent to the mean mistranslation rate). Results are shown for 10^4^ adaptive walks on the antibody-binding landscape at high expression (500 proteins per cell) and a population size of *N* = 10^8^.

A second test of our hypothesis is based on its prediction that some synonymous mutations come under positive selection only under mistranslation. To detect this selection pressure we consider what mistranslation rates would have been in genotypes that are traversed by an adaptive walk without mistranslation. We call these inferred mistranslation rates post hoc mistranslation rates. By comparing the true and the post hoc mistranslation rates, we want to find out whether genotypes encountered by adaptive walks with mistranslation have different mistranslation rates than genotypes encountered by adaptive walks without mistranslation.

Indeed, while many of the synonymous mutations that reach fixation during adaptive walks with mistranslation have very weak fitness effects (both beneficial and deleterious), some have large fitness effects (fig. 5B). Such mutations tend to appear after at least 10^3^ mutations have occurred. They increase fitness by switching to synonymous codons with lower mistranslation rates and therefore higher fitness. For a population of *N* = 10^8^ individuals and with high expression, every synonymous mutation with a fitness effect greater than 0.002 reduces mistranslation rates on average by 4%. Synonymous mutations with larger fitness effects also decrease mistranslation rates more strongly (Kendall’s *τ* = *−*0.579, *p* ≪ 10^−300^, *n* = 5789), a relation that is much weaker for nonsynonymous mutations (Kendall’s tau 5.44 *×* 10^−3^, *p* = 0.0225, *n* = 78021). Consequently, in adaptive walks with mistranslation, we observe a long-term decrease in mistranslation rates that is largely (but not entirely, see supplementary section 7) absent from post hoc mistranslation rates in walks without mistranslation (fig. 5C). A similar pattern also holds for the two toxin-antitoxin landscapes, except that there nonsynonymous mutations also decrease mistranslation rates. For example, in the toxin-antitoxin (E3) landscape, nonsynonymous mutations with larger fitness effects decrease mistranslation rates (Kendall’s *τ* = *−*0.0205, *p* = 3.00 *×* 10^−16^, *n* = 70359), although the association is an order of magnitude stronger for synonymous mutations (Kendall’s *τ* = *−*0.476, *p* = 2.40 *×* 10^−164^, *n* = 1511 both correlations *N* = 10^8^ and high expression). In the toxin-antitoxin (E2) landscape, nonsynonymous mutations decrease mistranslation rates weakly (Kendall’s *τ* = *−*8.47 *×* 10^−3^, *p* = 5.68 *×* 10^−4^, *n* = 73593), and the association is stronger for synonymous mutations (Kendall’s *τ* = *−*0.628, *p* = 1.27 *×* 10^−180^, *n* = 944 both correlations *N* = 10^8^ and high expression).

We also found evidence that the fixation of synonymous mutations is not only in itself adaptive, but that it can facilitate adaptation by increasing the supply of beneficial non-synonymous mutations. For example, in the antibody-binding landscape for populations of *N* = 10^4^ individuals and with mistranslation at low expression, synonymous mutations increased the average number of nonsynonymous genetic neighbours with higher fitness than the current genotype from 5.38 *×* 10^−2^ before fixation of the synonymous mutation to 1.99 *×* 10^−1^ afterwards (Wilcoxon test *z* = 17119.5, *p* = 2.37 *×* 10^−244^, *n* = 9566). This effect is particularly strong once populations have reached high fitness and the supply of beneficial nonsynonymous mutations has become limited. In adaptive walks after the first 10^4^ mutations, the average number of beneficial nonsynonymous mutations increases by 94.7 percent from 7.22 *×* 10^−3^ before the fixation of a synonymous mutation to 1.36 *×* 10^−1^ afterwards (Wilcoxon test *z* = 7903.5, *p* = 9.17 *×* 10^−201^, *n* = 8730). A similar pattern holds for the other landscapes and in adaptive walks without mistranslation (supplementary table S4). Consequently, adaptive walks that fix few synonymous mutations, such as walks at high population sizes and without mistranslation, experience less adaptive change at high fitnesses.

## Discussion

Because the phenotypic mutations caused by mistranslation are much more frequent than DNA mutations, they represent an important source of variation that may affect adaptive evolution despite its low heritability. To find out how mistranslation may affect evolution, we use experimentally measured mistranslation rates to study its effects on the topography of experimentally characterised adaptive landscapes, and on how populations evolve on such landscapes. We find that mistranslation broadly decreases the fitness of high fitness genotypes and increases the fitness of low fitness genotypes, leading to a flattening of the adaptive landscape. However, mistranslation does not affect all genotypes equally. It changes the fitness of some genotypes relative to others. The most dramatic effect of mistranslation is a large decrease in the number of mutations that are nearly neutral, i.e. mutations whose impact on fitness is so small that they are effectively invisible to selection (Ohta 1973). More than half of these mutations are synonymous. Our analysis shows that mistranslation can cause small fitness differences between synonymous mutations. Most of those fitness differences are orders of magnitude smaller than the differences caused by nonsynonymous mutations, but they are still large enough to become subject to selection in species with large effective population sizes (Sung et al. 2012).

Previous theoretical work (Whitehead et al. 2008; Wang and Zhang 2011; Rocabert et al. 2020) has established that nonheritable variation in fitness can sometimes favour the fixation of beneficial mutations. However, it is not clear how frequently this occurs. On the landscapes we study, mistranslation increases the fixation probability of roughly three to eight percent of beneficial mutations. The reason is that mistranslation results in the production of protein variants that are more beneficial than the variant genetically encoded by the mutant. This advantage comes at the cost of decreasing the fixation probability of many other beneficial mutations. However, considering experimental studies that demonstrate the fitness benefits of mistranslation in populations under stress (Javid et al. 2014; Ribas de Pouplana et al. 2014; Mohler and Ibba 2017; Samhita et al. 2021) we suggest that mistranslation may often help the fixation of beneficial mutations in nature. We also find that mistranslation renders a small fraction of beneficial mutations deleterious, or vice versa (approximately two percent both ways). Thus, mistranslation can open new paths to higher fitness, and close others.

Our results also suggest that mistranslation helps evolving populations reach high fitness regions in rugged landscapes that are difficult to navigate, but conveys little to no benefit on landscapes where high fitness genotypes are easily found. Specifically, in adaptive walks on the antibody-binding landscape, where reaching high fitness genotypes is more difficult than on the toxin-antitoxin landscapes, populations with mistranslation reach on average a slightly higher fitness than populations without mistranslation. These advantages are small in part because the mistranslation rates we use in our study come from a population of *E. coli* growing in a favourable environment (Mordret et al. 2019). However, mistranslation rates increase in stressful conditons (Netzer et al. 2009), and consequently the benefits we report here may be larger in nature (Samhita et al. 2021).

Our analysis also supports a prediction by (Rocabert et al. 2020) that nonheritable variation can accelerate adaptation not only at low fitness, but also at high fitness. We find two causes for this phenomenon. First, selection favours the reduction of mistranslation rates at high fitness, because mistranslation is typically detrimental for high fitness genotypes. This reduction occurs predominantly through synonymous mutations. This cause is consistent with the experimental observation that selection favours synonymous substitutions that decrease mistranslation rates (Akashi 1994; Drummond and Wilke 2008; Porceddu et al. 2013; Zaborske et al. 2014). Second, the fixation of synonymous mutations can open new paths to even higher fitness. Once a population reaches high fitness, some synonymous mutations can act as potentiating mutations (Blount et al. 2008), i.e. mutations that themselves have a small or no impact on fitness but that increase the supply of beneficial nonsynonymous mutations. Mistranslation increases the probability that synonymous mutations become fixed, and therefore makes the discovery of these beneficial nonsynonymous mutations easier.

As a previous theoretical study on mistranslation has also observed (Drummond and Wilke 2008), our results suggest that mistranslation can render the ratio of nonsynonymous to synonymous mutations (dN/dS) unreliable as an indicator of the absence of positive selection. More precisely, a large dN/dS ratio indicates directional selection only early during adaptive evolution, when a population’s fitness increases dramatically through nonsynonymous mutations. This is no longer the case later during adaptive evolution, when most genotypes in a population have attained high fitness. The reason is that mistranslation is deleterious for high fitness genotypes, and selection favours synonymous mutations that decrease mistranslation rates. Since the dN/dS ratio is designed to report selection on nonsynonymous mutations, it thus under-reports the extent of directional selection.

Like any other computational analysis, ours is limited by inevitable simplifying assumptions. First, we assume that without mistranslation, synonymous mutations are neutral. That is not necessarily the case, because synonymous DNA mutations can alter mRNA stability (Duan et al. 2003; Chamary and Hurst 2005; Kristofich et al. 2018) or ribosome binding (Eyre-Walker and Bulmer 1993; Knöppel et al. 2016). One consequence of this assumption is that we may overestimate by how much mistranslation reduces the number of nearly neutral mutations, because many synonymous mutations may not be neutral to begin with. Second, our analysis requires a large amount of data. The few available sufficiently large empirical landscapes may not be representative of protein adaptive landscapes in general. This limitation will be alleviated as more and larger landscapes will be characterised in the future.

Third, the effects of mistranslation are more complicated in nature than in our model. For example, they may be buffered by metabolism (Kacser and Burns 1981), changes in gene expression (Bratulic et al. 2015), preferential degradation of misfolded proteins (Mohler and Ibba 2017), and DNA mutations that increase protein stability (Bratulic et al. 2017; Zheng et al. 2021). Conversely, they may also be enhanced, for example through protein aggregation (Bucciantini et al. 2002; Drummond and Wilke 2008). In addition, the fitness of organisms is not equally sensitive to changes in the activity of protein-coding genes, with some genes being very robust to changes in activity, and others very sensitive (Keren et al. 2016). We simulate a case where fitness is very sensitive to changes in the activity of a protein, and the effects of mistranslation may be weaker for most genes. These complicating factors are an exciting research opportunity to develop more realistic models. Because the phenomena we study are difficult to observe directly by experiment, such data-driven models will remain essential to study how mistranslation changes the evolutionary dynamics on adaptive landscapes.

## Materials and Methods

### Simulating the fitness effects of mistranslation

Our simulations start from a given genotype, that is, a nucleotide sequence encoding the part of the focal protein for which fitness data on protein variants is available (Wu et al. 2016; Lite et al. 2020). Each codon in this nucleotide sequence has a given probability of being translated into either the cognate amino acid or into one of the 19 noncognate amino acids at a rate of mistranslation that is experimentally determined (Mordret et al. 2019). Consequently, every time a nucleotide sequence of length 3*L* is translated, there is a finite chance of producing one of 20*^L^* protein variants of length *L*, each with a unique polypeptide sequence.

We model the distribution of protein variants resulting from (mis)translation in a population of unicellular organisms as a multinomial distribution with *n* trials, where *n* is the number of proteins produced per cell (a detailed model description and mathematical expressions are available in supplementary section 3). The probability of producing a given protein variant is set by the empirical mistranslation rates (Mordret et al. 2019). To estimate the probability of producing a given protein variant, we assume that the mistranslation rate at each codon is independent from that at other codons, because experimental measurements suggest that most translation errors are due to codon-anticodon mispairing in the ribosome (Kramer and Farabaugh 2007; Mordret et al. 2019). In order to save computational resources, we ignore protein variants that are very rarely produced from a given genotype (probability of being produced by errors in translation is less than 10^−9^). To estimate the fitness distribution of a population of cells experiencing mistranslation errors, we assume that every one of the *n* protein copies produced per cell contribute equally to fitness, and that these fitness contributions are independent of one another. The fitness of a cell is then the average of all fitness contributions of the *n* proteins produced. These fitness contributions are given by the fitness of fitness equivalents of the protein variants in the adaptive landscapes. The expected fitness of a cell *E*(*f*) is therefore the average of the fitness contributions of the expected set of protein variants predicted by the multinomial distribution. Equivalently, the variance in fitness *V ar*(*f*) among cells within the population is a function of the variance in protein composition predicted by the multinomial distribution. These two estimates are sufficient for the remainder of our analysis.

### Estimating the fixation probability of novel mutations

Given an estimate for the mean and variance in fitness for both a wild-type genotype *wt* and a single-nucleotide DNA mutant *mt* of the wild-type, we here estimate the probability of fixation of the mutant genotype if it is introduced into a population that consists of only wild-type genotypes. To do so, we need three quantities. The first one is the selection coefficient *s*, which we define as the relative difference between the expected fitness values of the invading mutant *E*(*f_mt_*) and of the wild type *E*(*f_wt_*):

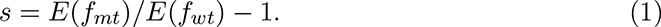

The second quantity is the effective population size *N_e_*. Mistranslation can decrease the power of selection by adding non-genetic variation in fitness. As a result, in a population under the influence of mistranslation the amount of variation in allele frequencies from one generation to the next is larger than in a population without mistranslation. Previous authors (Wang and Zhang 2011) have modelled this increase in the variation of allele frequencies between generations as being equivalent to a decrease in the effective population size of a haploid population from *N* to *N_e_*, according to

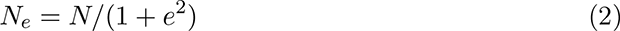

where *N* is the population size in the absence of mistranslation and *e*^2^ is the squared coefficient of variation in fitness (*V ar*(*f_wt_*)*/E*(*f_wt_*)^2^). We follow this approach. The new effective population size *N_e_* and the selection coefficient *s* can then be used to estimate the probability of fixation of the invading mutation, where the initial allele frequency is *p* = 1*/N*. (We note that the expression *p* = 1*/N_e_*would be incorrect here, because the initial allele frequency is not determined by the effective population size *N_e_* but by the actual number of individuals *N*). The third quantity is the fixation probability *u_fix_*. We use Kimura’s equation (Kimura 1962) to estimate the fixation probability of a new mutant in a haploid population:

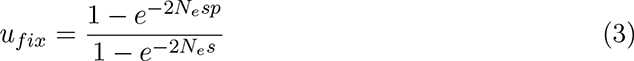

### Simulating adaptive walks on an adaptive landscape

We simulate adaptive evolution on our three landscapes at the level of nucleotide sequences. However, the landscape data is reported at the amino acid sequence level Wu et al. 2016; Lite et al. 2020. We therefore make the simplifying assumption that synonymous mutations are neutral in the absence of mistranslation.

We simulate the occurrence and either fixation or loss of mutations, and not the gradual change in allele frequency of a population from one generation to the next. Behind this procedure lies the assumption that the evolutionary dynamics on our landscapes fall into the weak mutation-strong selection regime (Gillespie 1983; Gillespie 1984). The reason is that we simulate evolution at a small number of nucleotide sites, where it is unlikely that multiple alleles will be segregating in a population at these sites at any one time.

During each step of our simulations, we choose a one-step mutational neighbour of the current (wild-type) genotype at random, and evaluate its probability of fixation. Based on this probability, the mutation either reaches fixation or is lost at random. We continue this process for 10^5^ successive mutations per walk, and note that only a small fraction of these mutations reach fixation. We perform 10^4^ replicate adaptive walks for each evolutionary scenario we model, each with its own starting genotype chosen at random from among the 10% of genotypes with lowest fitness.

## Acknowledgements

This project has received funding from the European Research Council (https://erc.europa.eu/) under Grant Agreement No. 739874. We would also like to acknowledge support by the Swiss National Science Foundation (http://www.snf.ch/en/Pages/default.aspx) grant 31003A_172887, by the University Priority Research Program in Evolutionary Biology (https://www.evolution.uzh.ch/en.html), and the Forschungskredit of the University of Zurich, grant no. FK-21-103. The funders had no role in study design, data collection and analysis, decision to publish, or preparation of the manuscript.

## Data availability

The simulation, data handling, and analysis code is publicly available from https://github.com/michaelacmschmutzer/lost_in_translation.

