## Supplementary materials for "Not quite lost in translation: Mistranslation alters adaptive landscape topography and the dynamics of evolution"

### Supplementary appendix S1 to “Not quite lost in translation: Mistranslation alters adaptive landscape topography and the dynamics of evolution”

**4 Mistranslation shrinks the size of nearly neutral networks 17**

**5 Mistranslation has small or inconsistent effects on the fitness reached by the end of adaptive walks on the toxin-antitoxin landscapes 20**

**6 Protein expression level influences adaptive walk trajectory 20**

**7 Post-hoc mistranslation rates can decrease in walks without mistranslation 21**

**8 Approximating mistranslation-free fitness 22**

### 1 Supplementary tables

| Landscape | $N$ | Condition | $\bar{x}$ (%) | $\sigma$ (%) | $H$ | $p$-value |
|---|---|---|---|---|---|---|
| | | No mistranslation | 50.2 | 27.7 | | |
| Antibody-binding | $10^4$ | Mistranslation 1 protein | 52.1 | 26.8 | 44.6 | $2.05 \times 10^{-10}$ |
| | | Mistranslation 500 proteins | 51.8 | 27.2 | | |
| | | No mistranslation | 93.4 | 12.8 | | |
| E3 toxin-antitoxin | $10^4$ | Mistranslation 1 protein | 93.9 | 12.0 | 184 | $9.06 \times 10^{-41}$ |
| | | Mistranslation 500 proteins | 93.6 | 12.7 | | |
| | | No mistranslation | 95.2 | 14.2 | | |
| E2 toxin-antitoxin | $10^4$ | Mistranslation 1 protein | 95.5 | 13.5 | 815 | $1.07 \times 10^{-177}$ |
| | | Mistranslation 500 proteins | 95.0 | 14.7 | | |
| | | No mistranslation | 47.6 | 27.2 | | |
| Antibody-binding | $10^6$ | Mistranslation 1 protein | 50.2 | 26.3 | 80 | $3.79 \times 10^{-18}$ |
| | | Mistranslation 500 proteins | 49.9 | 26.5 | | |
| | | No mistranslation | 92.0 | 13.4 | | |
| E3 toxin-antitoxin | $10^6$ | Mistranslation 1 protein | 92.4 | 13.1 | 134 | $7.59 \times 10^{-30}$ |
| | | Mistranslation 500 proteins | 92.5 | 13.3 | | |
| | | No mistranslation | 94.1 | 14.8 | | |
| E2 toxin-antitoxin | $10^6$ | Mistranslation 1 protein | 94.3 | 14.5 | 436 | $1.97 \times 10^{-95}$ |
| | | Mistranslation 500 proteins | 94.2 | 14.7 | | |
| | | No mistranslation | 47.3 | 27.2 | | |
| Antibody-binding | $10^8$ | Mistranslation 1 protein | 50.2 | 26.1 | 95.3 | $2.03 \times 10^{-21}$ |
| | | Mistranslation 500 proteins | 49.9 | 26.5 | | |
| | | No mistranslation | 92.0 | 13.4 | | |
| E3 toxin-antitoxin | $10^8$ | Mistranslation 1 protein | 92.3 | 13.3 | 151 | $1.73 \times 10^{-33}$ |
| | | Mistranslation 500 proteins | 92.5 | 12.9 | | |
| | | No mistranslation | 93.9 | 15.3 | | |
| E2 toxin-antitoxin | $10^8$ | Mistranslation 1 protein | 94.6 | 13.8 | 337 | $6.78 \times 10^{-74}$ |
| | | Mistranslation 500 proteins | 94.1 | 15.1 | | |

Table S1: Comparison of the fitness values at the end of adaptive walks on all three adaptive landscapes. Mean ($\bar{x}$) and standard deviation ($\sigma$) of these fitness values are reported as a percentage of the maximum fitness in each landscape. For each combination of landscapes and population sizes $N$, we perform Kruskal-Wallis tests to determine if any significant differences exist between adaptive walks under three conditions. These conditions are without mistranslation, with mistranslation and low expression (one protein per cell), or with mistranslation and high expression (500 proteins per cell). We report the resulting $H$ statistic and $p$-value. Each set of conditions has $10^4$ replicates. Pairwise comparisons using Dunn's post-hoc test with Bonferroni correction of the significance of differences between conditions are given in table S2.

| Landscape | $N$ | Condition 1 | Condition 2 | $p$-value |
|---|---|---|---|---|
| | | No mistranslation | Mistranslation 1 protein | $2.82 \times 10^{-9}$ |
| Antibody-binding | $10^4$ | No mistranslation | Mistranslation 500 proteins | $2.25 \times 10^{-7}$ |
| | | Mistranslation 1 protein | Mistranslation 500 proteins | 1.00 |
| | | No mistranslation | Mistranslation 1 protein | $4.29 \times 10^{-32}$ |
| E3 toxin-antitoxin | $10^4$ | No mistranslation | Mistranslation 500 proteins | $8.48 \times 10^{-31}$ |
| | | Mistranslation 1 protein | Mistranslation 500 proteins | 1.00 |
| | | No mistranslation | Mistranslation 1 protein | $1.54 \times 10^{-129}$ |
| E2 toxin-antitoxin | $10^4$ | No mistranslation | Mistranslation 500 proteins | $3.18 \times 10^{-139}$ |
| | | Mistranslation 1 protein | Mistranslation 500 proteins | 1.00 |
| | | No mistranslation | Mistranslation 1 protein | $2.06 \times 10^{-14}$ |
| Antibody-binding | $10^6$ | No mistranslation | Mistranslation 500 proteins | $3.32 \times 10^{-14}$ |
| | | Mistranslation 1 protein | Mistranslation 500 proteins | 1.00 |
| | | No mistranslation | Mistranslation 1 protein | $3.27 \times 10^{-21}$ |
| E3 toxin-antitoxin | $10^6$ | No mistranslation | Mistranslation 500 proteins | $5.21 \times 10^{-25}$ |
| | | Mistranslation 1 protein | Mistranslation 500 proteins | 1.00 |
| | | No mistranslation | Mistranslation 1 protein | $3.42 \times 10^{-72}$ |
| E2 toxin-antitoxin | $10^6$ | No mistranslation | Mistranslation 500 proteins | $4.48 \times 10^{-73}$ |
| | | Mistranslation 1 protein | Mistranslation 500 proteins | 1.00 |
| | | No mistranslation | Mistranslation 1 protein | $4.70 \times 10^{-17}$ |
| Antibody-binding | $10^8$ | No mistranslation | Mistranslation 500 proteins | $1.52 \times 10^{-16}$ |
| | | Mistranslation 1 protein | Mistranslation 500 proteins | 1.00 |
| | | No mistranslation | Mistranslation 1 protein | $9.06 \times 10^{-27}$ |
| E3 toxin-antitoxin | $10^8$ | No mistranslation | Mistranslation 500 proteins | $4.21 \times 10^{-25}$ |
| | | Mistranslation 1 protein | Mistranslation 500 proteins | 1.00 |
| | | No mistranslation | Mistranslation 1 protein | $8.13 \times 10^{-57}$ |
| E2 toxin-antitoxin | $10^8$ | No mistranslation | Mistranslation 500 proteins | $4.91 \times 10^{-56}$ |
| | | Mistranslation 1 protein | Mistranslation 500 proteins | 1.00 |

Table S2: Pairwise comparison of the significance of differences between the fitness values at the end of adaptive walks without mistranslation, with mistranslation and low expression (one protein per cell), and with mistranslation and high expression (500 proteins per cell). All comparison are done between walks on the same landscape at the same population size $N$, and with $10^4$ replicates. We use Dunn's test with Bonferroni correction to assess significance.

| Landscape | $N$ | Condition | $\bar{x}$ | $\sigma$ |
|---|---|---|---|---|
| | | No mistranslation | 7.7 | 3.1 |
| Antibody-binding | $10^8$ | Mistranslation 1 protein | 7.5 | 3.0 |
| | | Mistranslation 500 proteins | 8.4 | 3.1 |
| | | No mistranslation | 7.0 | 2.6 |
| E3 toxin-antitoxin | $10^8$ | Mistranslation 1 protein | 7.1 | 2.6 |
| | | Mistranslation 500 proteins | 7.2 | 2.6 |
| | | No mistranslation | 7.3 | 2.5 |
| E2 toxin-antitoxin | $10^8$ | Mistranslation 1 protein | 7.3 | 2.4 |
| | | Mistranslation 500 proteins | 7.5 | 2.5 |
| | | No mistranslation | 7.8 | 3.1 |
| Antibody-binding | $10^6$ | Mistranslation 1 protein | 7.5 | 3.0 |
| | | Mistranslation 500 proteins | 8.4 | 3.2 |
| | | No mistranslation | 7.0 | 2.6 |
| E3 toxin-antitoxin | $10^6$ | Mistranslation 1 protein | 7.1 | 2.6 |
| | | Mistranslation 500 proteins | 7.2 | 2.6 |
| | | No mistranslation | 7.4 | 2.4 |
| E2 toxin-antitoxin | $10^6$ | Mistranslation 1 protein | 7.3 | 2.4 |
| | | Mistranslation 500 proteins | 7.4 | 2.5 |
| | | No mistranslation | 10.5 | 3.7 |
| Antibody-binding | $10^4$ | Mistranslation 1 protein | 9.6 | 3.5 |
| | | Mistranslation 500 proteins | 10.5 | 3.6 |
| | | No mistranslation | 9.2 | 2.9 |
| E3 toxin-antitoxin | $10^4$ | Mistranslation 1 protein | 9.2 | 2.9 |
| | | Mistranslation 500 proteins | 9.2 | 3.0 |
| | | No mistranslation | 10.3 | 3.0 |
| E2 toxin-antitoxin | $10^4$ | Mistranslation 1 protein | 10.1 | 3.0 |
| | | Mistranslation 500 proteins | 10.3 | 3.1 |

Table S3: Number of mutations (mean $\bar{x}$ and standard deviation $\sigma$) that reach fixation during adaptive walks. Results are reported for three population sizes $N$ and three conditions: Adaptive walks without mistranslation, with mistranslation and low expression (one protein per cell), and with mistranslation and high expression (500 proteins per cell).

| Landscape | $N$ | Condition | $\bar{x}_{before}$ | $\sigma_{before}$ | $\bar{x}_{after}$ | $\sigma_{after}$ |
|---|---|---|---|---|---|---|
| | | No mistranslation | – | – | – | – |
| Antibody-binding | $10^8$ | Mistranslation 1 protein | $9.70 \times 10^{-2}$ | $4.64 \times 10^{-1}$ | $5.15 \times 10^{-1}$ | $5.95 \times 10^{-1}$ |
| | | Mistranslation 500 proteins | $8.02 \times 10^{-2}$ | $4.68 \times 10^{-1}$ | $4.52 \times 10^{-1}$ | $6.10 \times 10^{-1}$ |
| | | No mistranslation | – | – | – | – |
| E3 toxin-antitoxin | $10^8$ | Mistranslation 1 protein | $1.28 \times 10^{-1}$ | $3.52 \times 10^{-1}$ | $9.83 \times 10^{-1}$ | $3.15 \times 10^{-1}$ |
| | | Mistranslation 500 proteins | $5.95 \times 10^{-2}$ | $2.83 \times 10^{-1}$ | $9.05 \times 10^{-1}$ | $3.83 \times 10^{-1}$ |
| | | No mistranslation | – | – | – | – |
| E2 toxin-antitoxin | $10^8$ | Mistranslation 1 protein | $8.28 \times 10^{-1}$ | $1.00 \times 10^{0}$ | $1.52 \times 10^{0}$ | $6.88 \times 10^{-1}$ |
| | | Mistranslation 500 proteins | $7.14 \times 10^{-1}$ | $9.26 \times 10^{-1}$ | $1.34 \times 10^{0}$ | $6.84 \times 10^{-1}$ |
| | | No mistranslation | 0.00 | 0.00 | $1.11 \times 10^{0}$ | $3.33 \times 10^{-1}$ |
| Antibody-binding | $10^6$ | Mistranslation 1 protein | $2.29 \times 10^{-1}$ | $7.72 \times 10^{-1}$ | $5.68 \times 10^{-1}$ | $7.25 \times 10^{-1}$ |
| | | Mistranslation 500 proteins | $7.79 \times 10^{-2}$ | $4.22 \times 10^{-1}$ | $4.59 \times 10^{-1}$ | $5.61 \times 10^{-1}$ |
| | | No mistranslation | $4.76 \times 10^{-2}$ | $2.18 \times 10^{-1}$ | $1.05 \times 10^{0}$ | $3.84 \times 10^{-1}$ |
| E3 toxin-antitoxin | $10^6$ | Mistranslation 1 protein | $7.25 \times 10^{-2}$ | $2.97 \times 10^{-1}$ | $9.64 \times 10^{-1}$ | $3.73 \times 10^{-1}$ |
| | | Mistranslation 500 proteins | $8.29 \times 10^{-2}$ | $2.95 \times 10^{-1}$ | $9.64 \times 10^{-1}$ | $2.58 \times 10^{-1}$ |
| | | No mistranslation | $2.50 \times 10^{-1}$ | $6.22 \times 10^{-1}$ | $1.08 \times 10^{0}$ | $2.89 \times 10^{-1}$ |
| E2 toxin-antitoxin | $10^6$ | Mistranslation 1 protein | $6.00 \times 10^{-1}$ | $9.14 \times 10^{-1}$ | $1.34 \times 10^{0}$ | $6.39 \times 10^{-1}$ |
| | | Mistranslation 500 proteins | $5.35 \times 10^{-1}$ | $8.82 \times 10^{-1}$ | $1.51 \times 10^{0}$ | $5.51 \times 10^{-1}$ |
| | | No mistranslation | $2.68 \times 10^{-3}$ | $5.37 \times 10^{-2}$ | $1.17 \times 10^{-1}$ | $3.41 \times 10^{-1}$ |
| Antibody-binding | $10^4$ | Mistranslation 1 protein | $7.22 \times 10^{-3}$ | $1.20 \times 10^{-1}$ | $1.36 \times 10^{-1}$ | $3.86 \times 10^{-1}$ |
| | | Mistranslation 500 proteins | $4.89 \times 10^{-3}$ | $8.81 \times 10^{-2}$ | $1.35 \times 10^{-1}$ | $3.82 \times 10^{-1}$ |
| | | No mistranslation | $4.59 \times 10^{-2}$ | $2.35 \times 10^{-1}$ | $3.58 \times 10^{-1}$ | $5.77 \times 10^{-1}$ |
| E3 toxin-antitoxin | $10^4$ | Mistranslation 1 protein | $6.29 \times 10^{-2}$ | $3.06 \times 10^{-1}$ | $3.87 \times 10^{-1}$ | $6.13 \times 10^{-1}$ |
| | | Mistranslation 500 proteins | $6.41 \times 10^{-2}$ | $3.02 \times 10^{-1}$ | $3.80 \times 10^{-1}$ | $6.14 \times 10^{-1}$ |
| | | No mistranslation | $8.48 \times 10^{-2}$ | $2.81 \times 10^{-1}$ | $2.68 \times 10^{-1}$ | $5.62 \times 10^{-1}$ |
| E2 toxin-antitoxin | $10^4$ | Mistranslation 1 protein | $4.87 \times 10^{-2}$ | $2.83 \times 10^{-1}$ | $2.33 \times 10^{-1}$ | $5.20 \times 10^{-1}$ |
| | | Mistranslation 500 proteins | $5.48 \times 10^{-2}$ | $3.01 \times 10^{-1}$ | $2.38 \times 10^{-1}$ | $5.26 \times 10^{-1}$ |

Table S4: Number of potential beneficial nonsynonymous mutations before and after the fixation of synonymous mutations during adaptive walks after the first $10^4$ mutations (mean $\bar{x}$ and standard deviation $\sigma$). Dashes (–) indicate that no synonymous mutations occurred in the interval we examined.

#### 2 Supplementary figures

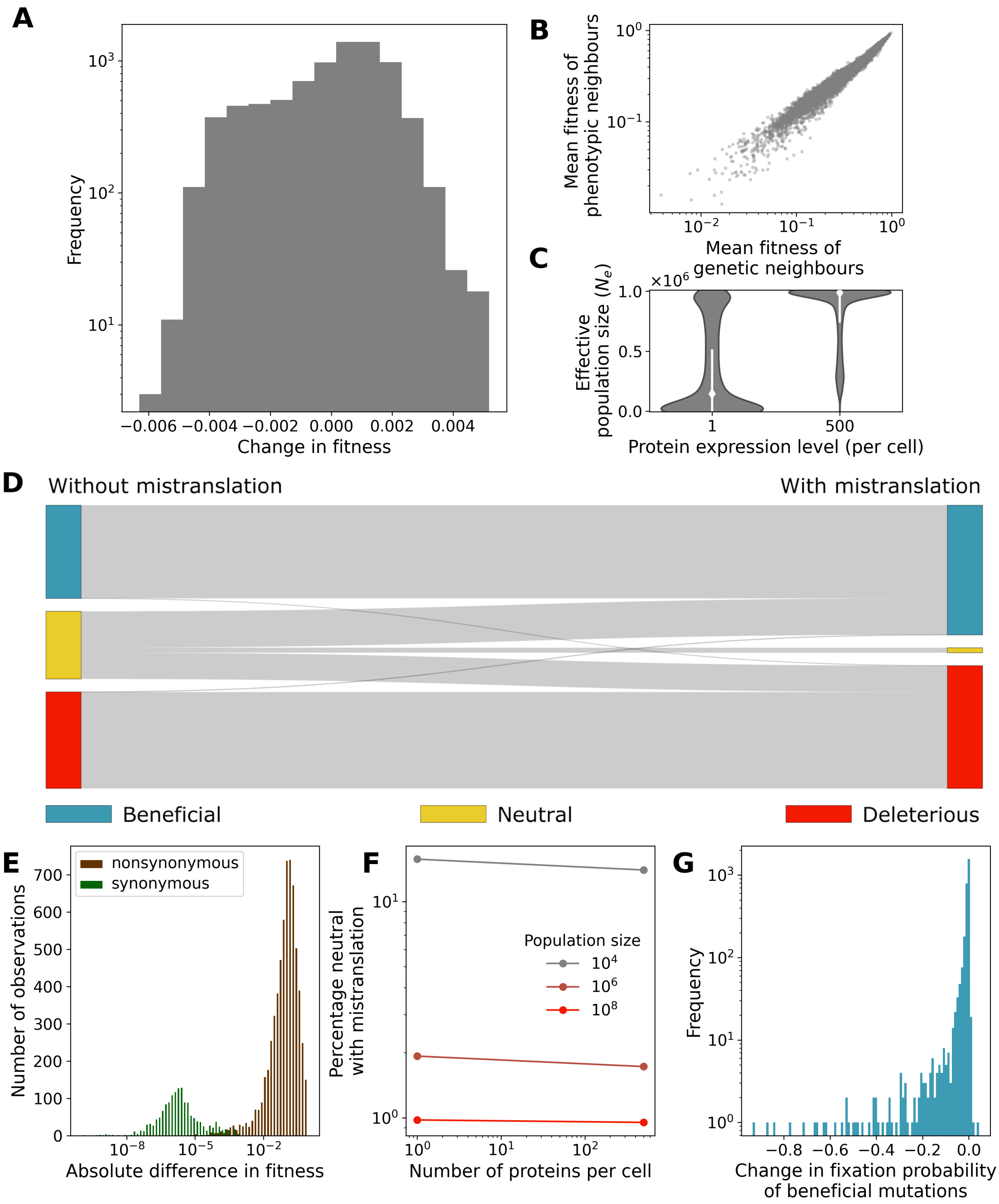

Figure S1: Mistranslation flattens the fitness distribution of the toxin-antitoxin (E3) landscape, and changes the fitness effects of some mutations. (caption continued next page)

Figure S1: (caption continued) A) Distribution of the changes in fitness due to mistranslation of $10^4$ randomly chosen genotypes. B) The mean fitness of a genotype's phenotypic neighbours (polypeptide sequences that differ in one amino acid position from the genotype, vertical axis) and the mean fitness of its genotypic neighbours (genotypes with a single nucleotide change, horizontal axis) are positively correlated. C) Mistranslation reduces the effective population size. White diamonds and lines show the median effective population size and the standard deviation, respectively. The violin plots show a Gaussian kernel density estimate of the distribution of effective population sizes. D) Changes due to mistranslation in the proportions of mutations classified according to their fitness effects as beneficial, neutral (including nearly neutral [1, 2]), or deleterious. E) Distributions of the absolute differences in mean fitness between both synonymous (green) and nonsynonymous (brown) pairs of genotypes sampled from the toxin-antitoxin (E3) landscape in the presence of mistranslation at an expression level of one protein per cell. Only nonzero fitness differences are shown. F) Percentage of mutations classified as neutral in the presence of mistranslation for multiple combinations of the population size $N$ and the protein expression level, both of which impact the effective population size $N_e$. G) Distribution of the change in fixation probabilities of beneficial mutations due to mistranslation. Data is shown only for those mutations that are beneficial both with and without mistranslation. The horizontal axis shows the difference in fixation probability due to mistranslation, with zero denoting no change. For example, changes in fixation probability close to negative one signify mutations that are almost certain to fix without mistranslation ($u_{fix} \approx 1$), but have almost no chance of reaching fixation with mistranslation ($u_{fix} \approx 0$). In all panels, results are shown for the same set of $10^4$ randomly chosen genotypes from the toxin-antitoxin (E3) landscape, and randomly chosen one-step mutational neighbours, at a population size of $N = 10^6$, and an expression level of one protein per cell.

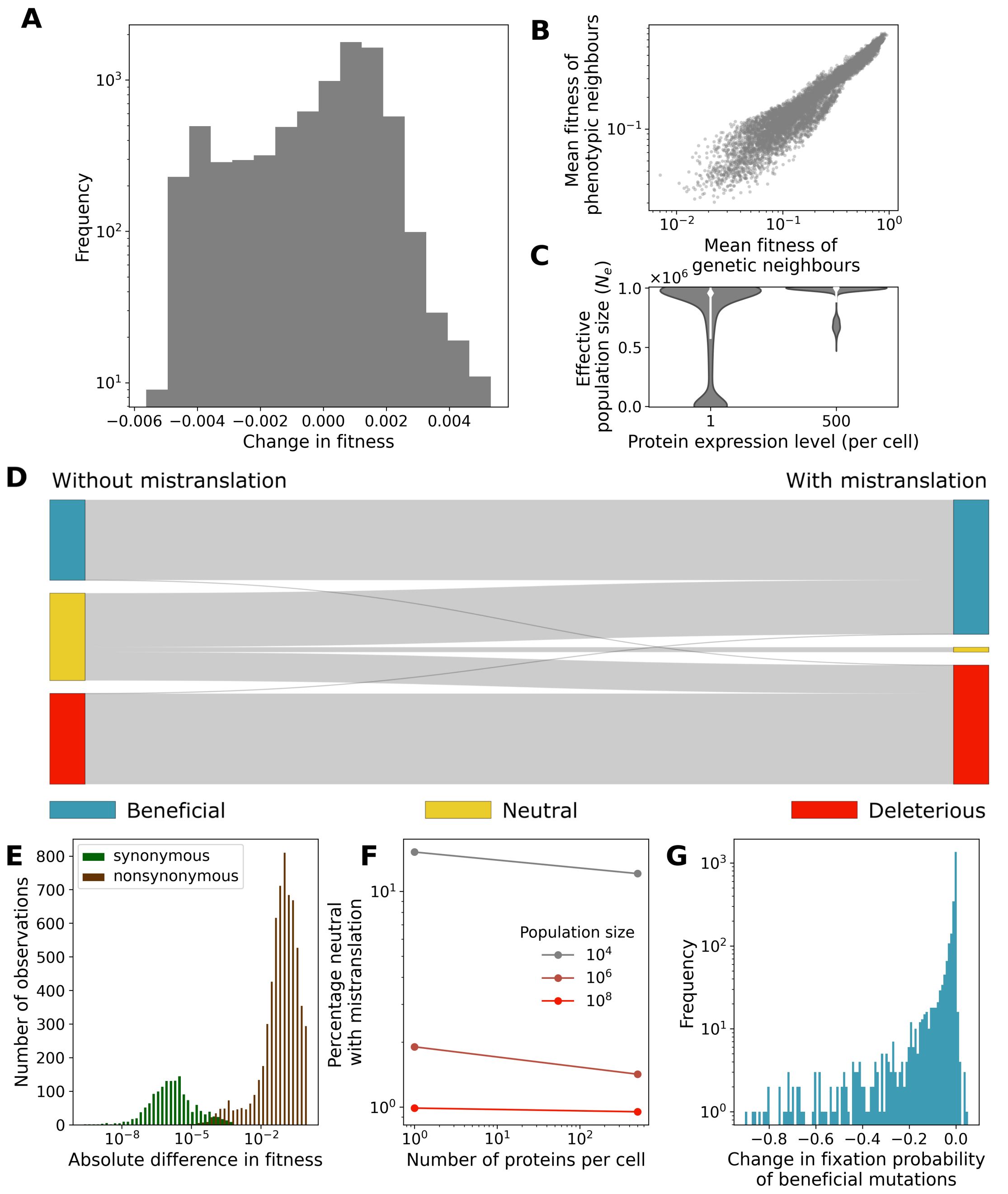

Figure S2: Mistranslation transforms the toxin-antitoxin (E2) landscape. (caption continued next page)

Figure S2: (caption continued) A) Distribution of the changes in fitness due to mistranslation of $10^4$ randomly chosen genotypes. B) The mean fitness of a genotype's phenotypic neighbours (polypeptide sequences that differ in one amino acid position from the genotype, vertical axis) and the mean fitness of its genotypic neighbours (genotypes with a single nucleotide change, horizontal axis) are positively correlated. C) Mistranslation reduces the effective population size. White diamonds and lines show the median effective population size and the standard deviation, respectively. The violin plots show a Gaussian kernel density estimate of the distribution of effective population sizes. D) Changes due to mistranslation in the proportions of mutations classified according to their fitness effects as beneficial, neutral (including nearly neutral [1, 2]), or deleterious. E) Distributions of the absolute differences in mean fitness between both synonymous (green) and nonsynonymous (brown) pairs of genotypes sampled from the toxin-antitoxin (E2) landscape in the presence of mistranslation at an expression level of one protein per cell. Only nonzero fitness differences are shown. F) Percentage of mutations classified as neutral in the presence of mistranslation for multiple combinations of the population size $N$ and the protein expression level, both of which impact the effective population size $N_e$. G) Distribution of the change in fixation probabilities of beneficial mutations due to mistranslation. Data is shown only for those mutations that are beneficial both with and without mistranslation. The horizontal axis shows the difference in fixation probability due to mistranslation, with zero denoting no change. For example, changes in fixation probability close to negative one signify mutations that are almost certain to fix without mistranslation ($u_{fix} \approx 1$), but have almost no chance of reaching fixation with mistranslation ($u_{fix} \approx 0$). In all panels, results are shown for the same set of $10^4$ randomly chosen genotypes from the toxin-antitoxin (E2) landscape, and randomly chosen one-step mutational neighbours, at a population size of $N = 10^6$, and an expression level of one protein per cell.

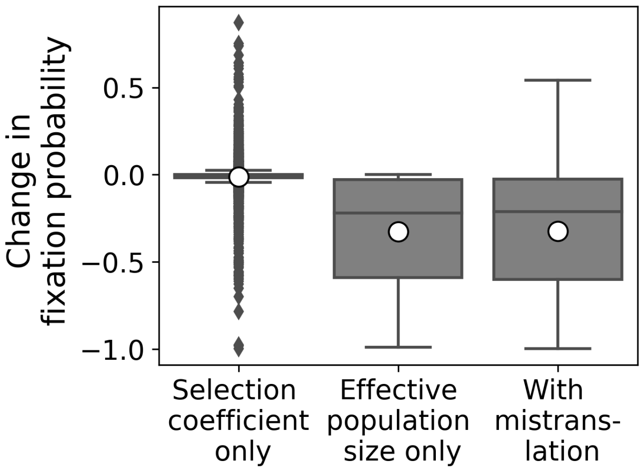

Figure S3: Changes in the effective population size dominate the effect of mistranslation on the fixation probability of beneficial mutations. Box-whisker plots of the distribution of changes in fixation probability of beneficial mutations when fixation probabilities are only affected by changes in selection coefficient caused by mistranslation (‘Selection coefficient only’), when only the effective population size changes (‘Effective population size only’), or when both mechanisms of mistranslation are taken into account (‘With mistranslation’). Boxes span the first to the third quantile of the distribution, and whiskers extend to 1.5 times the interquartile range from the box edges. Horizontal lines in each box denote the median, white circles show the mean of the distributions. Results are shown for 2813 genotypes from the antibody-binding landscape, and the same number of one-step mutational neighbours with higher fitness without mistranslation, at a populations size of $N = 10^6$, and an expression level of one protein per cell.

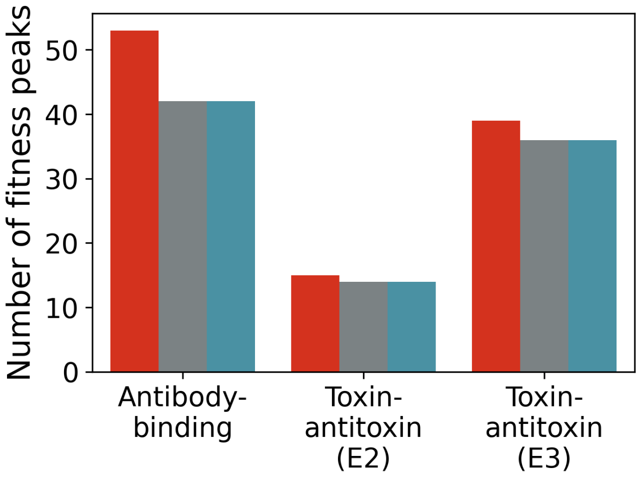

Figure S4: Mistranslation decreases the ruggedness of all three landscapes at low population sizes ($N = 10^4$). Number of fitness peaks among $10^5$ genotypes with the highest fitness in all three landscapes, without mistranslation (red) and with mistranslation at both high and low protein expression levels (500 and one protein per cell, grey and blue, respectively).

### 3 Supplementary materials and methods

#### 3.1 Model details

Our model of mistranslation begins with a nucleotide sequence encoding part of the focal protein, and the mistranslation rates of all the codons within the nucleotide sequence, i.e. the rates at which translation incorporates one of the noncognate amino-acids instead of the cognate amino acid. We assume that the probability of correct translation is one minus the sum of all mistranslation rates. For every polypeptide sequence of length $L$ encoded by a nucleotide sequence of length $3L$, (mis)translation produces from this nucleotide sequence one of $k = 20^L$ protein variants with a unique polypeptide sequence.

Any one cell that expresses the focal protein will harbour multiple variants of this protein, one resulting from correct translation, and all others from mistranslation. We estimate the number of each mistranslated variant in the total pool of $n$ copies of the focal protein in the cell through a multinomial distribution with $n$ 'trials'. Each trial corresponds to the biosynthesis of a protein, and can have one of $k$ possible outcomes – a protein variant. Each protein variant $i$ (with $i = 1, \ldots, k$) has its own probability of being produced $p_i$, which is determined by the mistranslation rates of the encoding nucleotide sequence. We assume that the mistranslation rate at each codon is independent from that at other codons, because experimental measurements suggest that most translation errors are due to codon-anticodon mispairing in the ribosome [3, 4]. Consequently, the probability $p_i$ of producing a given protein variant $i$ is the product of the probabilities that translation incorporates the amino acids that make up the protein's sequence, given the encoding nucleotide sequence. More specifically, the multinomial distribution for the number of copies $x_i$

of each protein variant $i,\dots,k$ is given by

$$f(x_1,\dots,x_k) = \begin{cases} \frac{n!}{x_1!\dots x_k!} p_1^{x_1} \times \dots \times p_k^{x_k}, & \text{if } \sum_{i=1}^{k} x_i = n \\ 0, & \text{otherwise.} \end{cases} \quad (1)$$

A single cell producing $n$ copies of the focal protein will contain a tiny fraction of all possible protein variants $k$. However, large populations of such cells will produce a larger fraction of variants. Over multiple generations or evolutionary time-scales, even rare protein variants will appear. Consequently, for the purpose of deriving our model we assume that all protein variants $k$ may be produced from a given nucleotide sequence. In practice, some of these variants will have such low probability of being produced through mistranslation that they have no influence on the fitness or evolution of a population, and in our simulations we ignore these variants (see below).

With this multinomial distribution we then calculate the expected number of copies of each protein variant, together with the variation in the number of copies of a given variant and the covariation in the number of copies between different protein variants. We need all three quantities to estimate the fitness distribution of a population of cells under the influence of mistranslation. The expected number of copies $E(x_i)$ of protein variant $i$ produced in a cell is

$$E(x_i) = np_i \quad (2)$$

and its variance is

$$Var(x_i) = np_i(1-p_i). \quad (3)$$

We calculate these two quantities for the number of copies of each possible protein variant $x_1,\dots,x_k$. Because the number of proteins produced per cell is finite, the production of a copy of one protein variant decreases the probability that a cell will contain a copy of another protein variant. Consequently, the covariance between the number of copies of different variants is negative. Specifically, the covariance of the number of copies $x_i$ of variant $i$ with the number of copies $x_j$ of another variant $j$ is

$$Cov(x_i,x_j) = -np_ip_j,\ i \neq j. \quad (4)$$

Using the multinomial distribution of a cell's protein composition together with experimental fitness measurements we then determine the fitness distribution $f$ of a population of cells expressing the protein. Specifically, we aim to estimate the mean and variance of the fitness distribution, because these quantities are needed later to estimate the fixation probability of mutations. The fitness of an individual cell depends upon the kind of protein variants it produces. For simplicity, we assume that every protein copy contributes equally to fitness, and that the fitness contribution $w_i$ of copy $i$ is independent of that of all other copies produced. The fitness of a cell then becomes the mean of the fitness contributions of all copies it produces. In consequence, the expected fitness of a cell $E(f)$ is given by

$$E(f) = \frac{1}{n}\sum_{i=1}^{k} n \cdot p_i \cdot w_i \quad (5)$$

where $n$ is the total number of proteins per cell, $k$ the number of all possible protein variants, $p_i$ the probability of translation producing protein variant $i$, and $w_i$ the fitness contribution of that protein variant. As indicated in this expression for $E(f)$, we calculate the expected (mean) fitness of a cell by summing over all the fitness contributions $w_i$ of each possible protein variant ($k$ in total), with variants that are more frequent having a greater impact, and then take the average by dividing this sum by the number of copies per cell $n$.

To arrive at an expression for the variance in fitness among cells, we start by observing in equation 5 that the expression for the expected fitness of a cell is a sum over the expected number of protein copies $n \cdot p_i$ for each protein variant $i$, multiplied by the term $w_i/n$. With this observation in mind, we compute the fitness variance by drawing on three well-known statistical principles. First, the variance of a random variable multiplied by a constant $a$ is $Var(a \cdot X) = a^2 \cdot Var(X)$. Consequently, every protein variant $i$ contributes to the variance in fitness according to the expression $Var(w_i/n \cdot x_i) = (w_i/n)^2 \cdot np_i(1-p_i)$. Second, the covariance of two random variables multiplied by the constants $a$ and $b$ is given by $Cov(a \cdot X, b \cdot Y) = a \cdot b \cdot Cov(X,Y)$. Consequently, the covariance in the fitness contributions of two protein variants $i$ and $j$ becomes $Cov(w_i/n \cdot x_i, w_j/n \cdot x_j) = -n \cdot p_i \cdot p_j \cdot w_i/n \cdot w_j/n$. Third, the variance of the sum of two random variables $X$ and $Y$ is $Var(X+Y) = Var(X) + Var(Y) + 2Cov(X,Y)$. Consequently, we can sum over the variances of the fitness contributions of all $k$ protein variants and adjust the resulting quantity by their pairwise (negative) covariances. Putting all these considerations together, the expected variance in the fitness of cells becomes

$$Var(f) = \frac{1}{n} \sum_{i=1}^{k} \left[ p_i \cdot (1-p_i) \cdot w_i^2 - \sum_{j=i+1}^{k} 2 \cdot p_i \cdot p_j \cdot w_i \cdot w_j \right]. \tag{6}$$

During our simulations we save computational resources by ignoring the contributions of all protein variants whose probability of being produced by translation was less than $10^{-9}$ when computing the mean fitness of cells and its variance.

#### 3.2 Mistranslation rate estimates

Following the literature in the field [5–7], we define a phenotypic mutation as the mistranslation of a specific codon into a specific (wrong) amino acid. We use experimentally estimated mistranslation rates from a previous study relying on proteome-wide mass-spectrometry in *E. coli* [4]. The study computed a ratio of mistranslated to correctly translated proteins. This ‘intensity ratio’ provides a direct estimate of the mistranslation rate per codon, which is reliable for common (but not for rare) mistranslation errors [4].

As an estimate for the rate at which any one phenotypic mutation occurs, we use the median intensity ratios of each phenotypic mutation for the wild-type *E. coli* strain growing in MOPS (3-(N-morpholino)propanesulfonic acid) complete medium during mid-exponential growth. As one might expect, these rate estimates decrease with the number of times any one phenotypic mutation is observed, except for rare phenotypic mutations, where mistranslation rate estimates *increase* as events become more rare, probably due to sampling error (supplementary figure S5a). Because direct rate estimates are therefore unreliable for rare phenotypic mutations, we do not use such estimates in our analysis. Instead, we estimate the rates of rare events through linear regression on the relationship between the number of times a phenotypic mutation was observed and the $\log_{10}$-transformed mistranslation rate estimates of common phenotypic mutations (supplementary figure S5a). After investigating multiple thresholds for the exclusion of direct rate estimates, we choose to exclude all mistranslation rate estimates with fewer than 18 observations, because this threshold minimises the intercept of the linear regression (supplementary figure S5b), and therefore minimises our estimate of the mistranslation rate for phenotypic mutations too rare to be observed in the empirical data [4]. This threshold excludes 86% of phenotypic mutations, for which we use indirect regression estimates. We assign unobserved phenotypic mutations the rate of $2.7 \times 10^{-4}$ per translated codon, the intercept of the regression line (supplementary figure S5a).

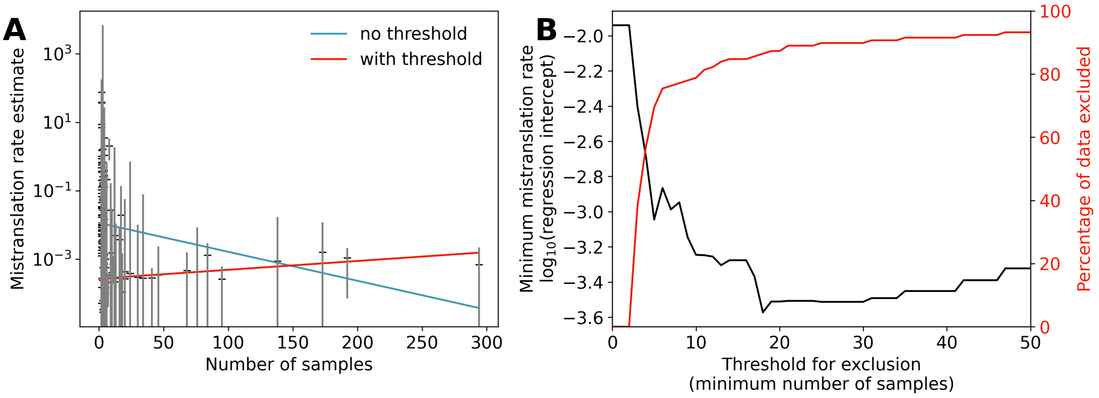

Figure S5: Interpolation of mistranslation rates using data from [4]. A) Correlation of the experimental mistranslation rate estimates of a given phenotypic mutation (vertical axis) with how many times the pertinent phenotypic mutation is observed (horizontal axis, each sample is an observation). Black crosses indicate median mistranslation rate estimates, and grey lines the corresponding 95 percent confidence intervals. The blue line shows the best linear regression fit when using all data, and the red line when all mistranslation rate estimates with less than 18 samples are ignored. B) Effect of different thresholds for exclusion (all mistranslation rate estimates with fewer samples than the threshold are ignored) on the intercept (black line) of the linear regression of the number of samples with mistranslation rate estimates (lines in panel A), and on the percentage of observations excluded from the regression analysis.

##### 3.3 Landscape data

We draw on three experimentally determined landscapes for all our analyses, as described in more detail below. The first of these is an antibody-binding landscape [8]. The second and third are toxin-antitoxin fitness landscapes that quantify the fitness of *E. coli* cells in which variants of an antitoxin bind to a toxin protein [9]. All landscapes are represented at the level of amino acids, because the nucleotide sequences encoding these amino acids have not been reported [8, 9]. We simulate evolution at the level of nucleotide sequences, so we map each nucleotide sequence to the amino acid it encodes. We chose these landscapes for two reasons. First, they contain phenotypic data for all 20 amino acids at each of the protein sites for which amino acid variation was examined. Second, they also contain data for all possible combinations of the 20 amino acids. In other words, these landscapes are combinatorially complete – they comprise data on all $20^L$ genotypes, where $L$ is the number of protein sites [8]. Because mistranslation can cause phenotypic mutations from any amino acid to any other amino acid [4], both characteristics are important for our purpose. We note that mistranslation inevitably affects these experimentally determined landscapes. Although it is partly possible to correct for this influence (supplementary section 9), we work with the empirical data, because (i) we are primarily interested in the relative changes induced by mistranslation, (ii) with the current data (only amino acid sequences are reported for the landscapes we study) we can only create rough approximations of the mistranslation-free fitness, and (iii) these approximations make predictions of the effect of mistranslation that are very similar to the predictions of our model (supplementary section 9).

The antibody-binding landscape [8] comprises *in vitro* data on the affinity of sequence variants of the protein GB1 to bind the antibody IgG-Fc. Specifically, it contains such data for all possible 20 amino acid variants at four sites in GB1, which is a streptococcal protein capable of binding immunoglobulins. Briefly, the authors of the pertinent study [8] created a mutant library encoding nearly all 160,000 GB1 sequence variants at the four amino acid positions, translated the library, and tagged the resulting proteins with the mRNA encoding them in order to identify the proteins later. They then incubated these tagged proteins with immunoglobulin-bearing beads, and washed away all non-bound proteins. They released the bound proteins and sequenced their mRNA tags. By comparing the frequencies of mRNA tags before and after exposure to immunoglobulin, they determined the binding activity of the GB1 mutants relative to the wild-type sequence. While this landscape is strictly speaking no fitness landscape, we follow other authors that have used a phenotype such as molecular binding data as proxies for fitness [10–13]. We thus tacitly assume that natural selection can favour strong molecular binding. The 160,000 amino acid sequences are encoded by more than $1.3 \times 10^7$ nucleotide sequences.

The two toxin-antitoxin landscapes [9] comprise fitness measures for almost 8,000 variants of the ParD3 antitoxin, a protein that is co-expressed with the toxin ParE3, which it binds and inhibits. To generate this data, amino acids at each of three sites in ParD3 were systematically replaced with all 19 other amino acids, and the resulting sequences were co-expressed in *E. coli* either with ParE3 (landscape 1) or its close homolog ParE2 (landscape 2 [9]). The fitness of *E. coli* cells expressing these variants was measured in mass selection experiments combined with deep sequencing, by measuring the change in frequency of each variant before and after selection for toxin-antitoxin binding [9]. Fitness values were transformed such that the most fit ParD3 variant was assigned a fitness of one, and variants of ParD3 with a nonsense mutation were assigned a fitness of zero. This transformation caused 8% and 17% of variants in the of the toxin-antitoxin (E3) and the toxin-antitoxin (E2) landscape, respectively, to have a fitness below zero. For the purpose of our analysis, we set all negative fitness values to zero, reasoning that at least some of them may have resulted from measurement error. The 8,000 amino acid sequences are encoded by about $2.2 \times 10^5$ nucleotide sequences.

#### 3.4 Counting the number of fitness peaks and the incidence of epistasis

In our analysis, a fitness peak can either consist of a single genotype or a network of multiple genotypes that are connected by single mutations, and in which genotypes that differ in a single mutation are nearly neutral [2], i.e. mutations with such small fitness effects that they effectively evolve by genetic drift. We define a nearly neutral mutation as a mutation with a selection coefficient $|s| < \frac{1}{4N_e}$ relative to the wild-type [1]. We consider such a genotype or network of genotypes a peak if all its single-mutation, non-neutral neighbours have lower fitness.

To search for fitness peaks in our three landscapes, we choose a set of $10^5$ genotypes with the highest fitness as seeds for the search. For each seed, we examine whether every one of the one-step mutational neighbours of the seed is nearly neutral or not. We test for neutrality both going from one genotype to another and in reverse, as the effective population size and selection coefficient can differ depending on the starting genotype. The reason is that genotypes may differ in the extent of between-individual fitness variation caused by mistranslation. We only consider a neighbouring genotype nearly neutral if mutations in both directions are nearly neutral. Sequences that match this criterion are members of the same nearly neutral network of genotypes. We then assess the near neutrality of *their* neighbours, repeating this process until we can not find any further nearly neutral neighbours. If one of the genotypes we chose as seeds is found to be a member of a previously explored nearly neutral network we continue on to another seed. During each step of this iterative search, we also keep track of the current nearly neutral network's non-neutral mutational neighbours. If all the non-neutral neighbours of each genotype in the network have a lower fitness than the genotype itself, we consider the set of genotypes in the network to constitute a fitness peak.

We estimate the incidence of epistasis by sampling $10^4$ genotypes from each of our landscapes. For each of these starting genotypes, we create a 'square' of four genotypes (see also supplementary figure S6). To do so, we choose two sites at random in the starting genotype (which we denote as 00) and replace either one or both sites with alternative nucleotides, generating an additional three nucleotide sequences for each starting genotype. Two of these sequences are single mutants (01 and 10), and one a double mutant (11). We quantify any deviation from additivity by using the linear combination of the fitness $w$ of each genotype [14]:

$$\varepsilon = w_{00} + w_{11} - w_{01} - w_{10}. \quad (7)$$

Any square that deviates from additivity ($\varepsilon \neq 0$) is epistatic. To differentiate between magnitude epistasis, simple sign epistasis, or reciprocal epistasis, we assess whether a given mutation causes the same sign change (positive or negative) in fitness in different genetic backgrounds, specifically the starting genotype (00) and the single mutant for the other site. If the background does not affect the sign change of neither mutation, we classify the square as a case of magnitude epistasis. If the background causes a sign reversal of one or both mutations, we classify the square as a case of simple sign epistasis or reciprocal sign epistasis, respectively.

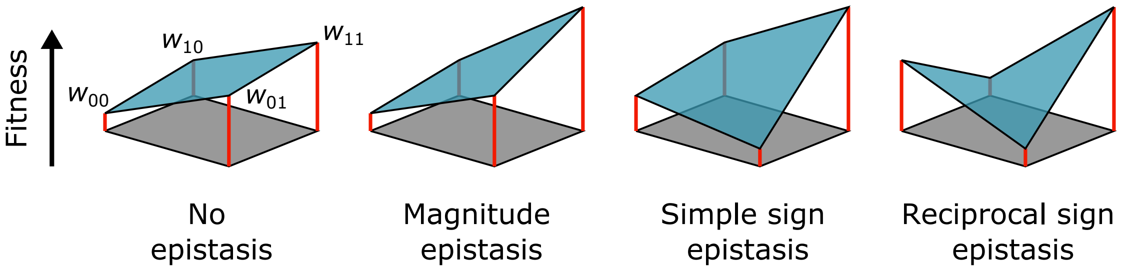

Figure S6: Fitness of four genotypes in either the absence of epistasis or under the influence of various kinds of epistasis. Two of the four genotypes are double mutants of each other (represented as 00 and 11), and two are single mutant intermediates (01 and 10). Together, these four genotypes form a 'square' (represented by the grey square on the bottom of each diagram, each genotype lies at one corner). Each genotype maps to a given fitness ($w$, height of red lines). Epistasis changes the fitness effects of mutations in different genetic backgrounds, causing genotypes to have a different relative fitness (blue shapes) from one another than would be expected in the absence of epistasis. Figure adapted from [15].

#### 4 Mistranslation shrinks the size of nearly neutral networks

In the main text we observe that mistranslation greatly reduces the number of nearly neutral mutations. This means that mistranslation is likely to disrupt networks of nearly neutral mutants. Here we quantify the extent of this disruption.

In order to find nearly neutral networks, we use the same method we use to find nearly neutral networks that are fitness peaks, except that we now consider such networks throughout the fitness landscape. We sample the distribution of nearly neutral network sizes by choosing a starting genotype and identifying all its nearly neutral mutational neighbours. We repeat this procedure with these neighbours, and so on, until we could not find any further nearly neutral neighbours. For the large antibody-binding landscape, we repeat this procedure for $10^4$ starting genotypes chosen at random, with the condition that each genotype should encode a unique amino acid sequence and have a fitness greater than zero. Because the smaller toxin-antitoxin landscapes have fewer than $10^4$ unique amino acid sequences, we instead choose one nucleotide sequence for each amino acid sequence with a fitness greater than zero as a starting genotype for this procedure, resulting in 7248 starting genotypes for the toxin-antitoxin (E3) landscape and 6533 genotypes for the toxin-antitoxin (E2) landscape. We determine the size distribution of the resulting neutral networks at three effective population sizes ($10^4$, $10^6$, $10^8$) in the presence or absence of mistranslation. We also explore the effect of protein expression level at either one or 500 proteins per cell. The former is an extreme value that can cause large reductions in effective population size, the latter is the median expression level in *E. coli* and is more broadly representative of expression levels in bacteria [16]. We use the same starting genotypes for all population sizes and expression levels.

In the absence of mistranslation, the size distribution of the sampled nearly neutral networks does not differ between the three population sizes we investigate, because most of the fitness differences between nonsynonymous neighbouring genotypes are sufficiently large to come under selection at the three effective population sizes we investigate (main text figure 3b). In the presence of mistranslation and at population sizes larger than $10^4$ we find very few networks for all landscapes. For example, for the toxin-antitoxin (E3) landscape at a population size of $10^6$ and an expression level of one protein per cell, only 21% of starting genotypes have at least one nearly neutral neighbour. The reason is that in the presence of mistranslation and in large populations, selection is sufficiently strong to act on the fitness differences between our starting genotypes and most of their mutational neighbours, both synonymous and nonsynonymous (figure S7). We will therefore describe our findings for the smallest population size $10^4$ only. In addition, we will focus on the results with a single protein per cell because the influence that mistranslation has on the size of neutral networks through changes in the effective population size is relatively small. By focusing on the extreme case of very low protein expression we quantify the effect that changes in effective population size can maximally have on the size of nearly neutral networks.

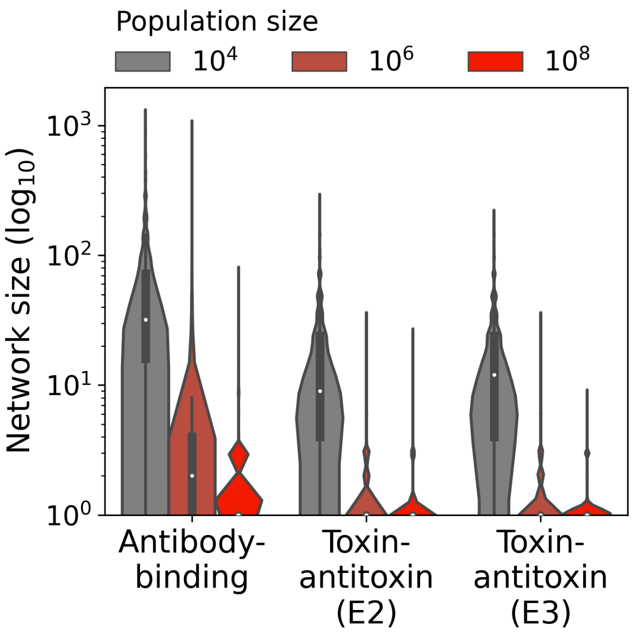

Figure S7: At large population sizes, mistranslation strongly decreases the size of neutral networks. The violin plots show Gaussian kernel density estimates of the distribution of network sizes. The boxes inside each violin plot span the first to the third quantile of the distribution, and vertical lines extend to 1.5 times the interquartile range from the box edges. White circles denote the median. The violin plots are shown only to a minimum neutral network size of one genotype, because values below that are not biologically meaningful. We determine all network sizes under mistranslation and under a protein expression level of one copy per cell.

Mistranslation causes a reduction by about one third in the average size of nearly neutral networks for all three landscapes (figure S8a). We expect that changes in the mean fitness of genotypes due to mistranslation decrease neutral network size, and changes in the effective population size increase neutral network size. In order to disentangle the contributions of these two conflicting effects, we repeat our analysis for the antibody-binding landscape, but ignore one of the two mechanisms by which mistranslation affects evolution (figure S8b). We find that reductions in effective population size due to mistranslation causes a 1% increase in the mean network size when we ignore changes in the mean fitness. In contrast, mistranslation's effect on a genotype's mean fitness leads to a one-half decrease in the mean size of nearly neutral networks when we ignore changes in effective population size. This reduction is two times as large as when we take both effects of mistranslation into account. Consequently, the effect of mistranslation on the size of nearly neutral networks is mediated by both changes in mean fitness and changes in the effective population size, and these two interact. This interaction is a consequence of the fact that the average fitness differences between nonsynonymous mutations are orders of magnitude larger than the average fitness differences between synonymous mutations caused by mistranslation (main text figure 3b). Reductions in the effective population size due to mistranslation are much more likely to overcome the smaller differences in mean fitness between synonymous mutations caused by mistranslation, and consequently changes in the effective population size have a larger impact on the size of neutral networks when we also take mistranslation's effect on mean fitness into account.

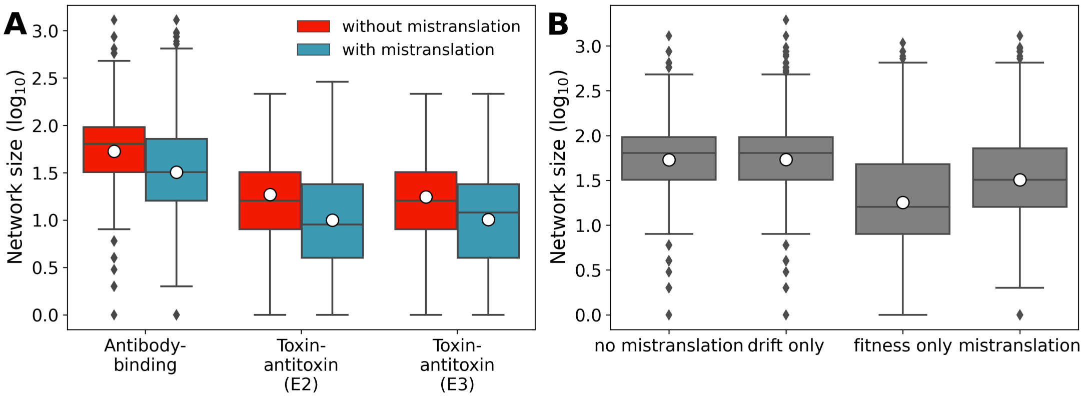

Figure S8: Mistranslation decreases nearly neutral network size, an effect that is mediated both by changes in the mean fitness of genotypes and changes in the effective population size $N_e$. A) Box-whisker plots of the distribution of nearly neutral network sizes in the presence (blue) and absence (red) of mistranslation for all three fitness landscapes. B) Box-whisker plots of the distribution of nearly neutral network sizes for networks in the antibody-binding landscape without mistranslation ('no mistranslation'), with mistranslation affecting only the effective population size $N_e$ ('drift only'), with mistranslation affecting only the mean fitness of genotypes ('fitness only'), and with mistranslation affecting both the effective population size and the mean fitness ('mistranslation'). Boxes span the first to the third quantile of the distribution, and whiskers extend to 1.5 times the interquartile range from the box edges. Horizontal lines in each box denote the median, white circles show the mean of the distributions. The protein expression level is one copy per cell and the population size is $N = 10^4$ in both panels.

### 5 Mistranslation has small or inconsistent effects on the fitness reached by the end of adaptive walks on the toxin-antitoxin landscapes

Mistranslation results in very small differences in the fitness reached by populations evolving on the toxin-antitoxin (E3) landscape. Specifically, walks without mistranslation reach 92.0± 13.4 percent of the maximum fitness and walks with mistranslation 92.3± 13.3 percent of the maximum. These differences are weak but statistically significant (Kruskal-Wallis test, $H = 151$, $p = 1.73 \times 10^{-33}$). Differences between walks with and without mistranslation are also significant (Dunn's test with Bonferroni correction, $p \leq 4.21 \times 10^{-27}$). In contrast, the effect of mistranslation on walks on the toxin-antitoxin (E2) landscape is inconsistent between different population sizes, with mistranslation being slightly beneficial at high population sizes but slightly detrimental in small populations. For example, at $N = 10^4$, populations without mistranslation reach on average 0.3 percent higher fitness than populations with mistranslation, i.e., 95.2±14.2 and 94.9±14.7 percent of the maximum fitness, respectively (Kruskal-Wallis test, $H = 815$, $p = 1.07 \times 10^{-177}$). Differences between walks with and without mistranslation are also significant (Dunn's test with Bonferroni correction, $p \leq 1.54 \times 10^{-129}$).

### 6 Protein expression level influences adaptive walk trajectory

Protein expression level can affect the trajectory of adaptive walks on the antibody binding landscape. The reason is that expression level modifies the effect of mistranslation on effective population size – at lower expression, between-individual variation in fitness is larger and the effective population size smaller. At the beginning of adaptive evolution the fitness of populations with mistranslation and high expression increases on average earlier than for populations with low expression (populations without mistranslation show an intermediate fitness increase). For example, 50 mutations after the beginning of the walk on the antibody-binding landscape ($N = 10^4$ individuals), populations without mistranslation have attained 21.4±21.4 percent of the maximum fitness, and populations with high and low expression have attained 22.9±21.7 percent and 19.0 (±20.8) percent of this maximum, respectively. The differences in fitness attained between populations with mistranslation and without, but also between different expression levels are statistically significant (Kruskal-Wallis test, $H = 309$, $p = 1.02 \times 10^{-67}$). All pairwise comparisons are statistically significant (Dunn's test with Bonferroni correction, $p \leq 2.37 \times 10^{-7}$). No such differences exist for the toxin-antitoxin landscapes. In summary, mistranslation can facilitate escape from low fitness genotypes in the antibody-binding landscape, and this process is faster in populations with high expression. The difference between adaptive walks with high and low expression shows that mistranslation is a double-edged sword. Mistranslation can accelerate adaptation, as long as it induces little nongenetic fitness variation between individuals. If it induces a lot of such variation, mistranslation can slow down adaptation.

#### 7 Post-hoc mistranslation rates can decrease in walks without mistranslation

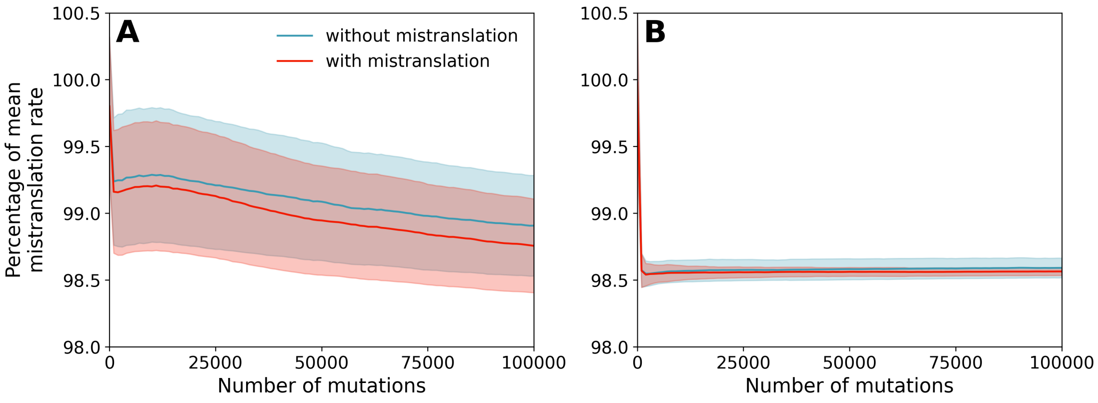

Figure S9: The structure of adaptive landscapes can cause adaptive walks to show systematic changes in post hoc mistranslation rates even in walks without mistranslation. A) Mean and standard error (lines and shaded areas, respectively) of the mistranslation rates (red) and post hoc mistranslation rates (blue) of genotypes encountered during adaptive walks on the toxin-antitoxin (E3) landscape. B) Mean and standard error (lines and shaded areas, respectively) of the mistranslation rates (red) and post hoc mistranslation rates (blue) of genotypes encountered during adaptive walks on the toxin-antitoxin (E2) landscape. For both panels, (post hoc) mistranslation rates are shown as percentages of the means of the mistranslation rates of all genotypes in their respective landscapes, where 100 percent stands for the mean mistranslation rate. Vertical axes show the number of mutations that have occurred since the beginning of the adaptive walk. Results are shown for $10^4$ walks at a population size of $10^4$ and, for walks with mistranslation, at a high expression level (500 proteins per cell).

We observe a slow reduction in post hoc mistranslation rates late during long adaptive walks without mistranslation (after the first $10^4$ mutations). At the end of such walks, populations have evolved high fitness, where mutations that increase fitness will also tend to decrease post hoc mistranslation rates. This reduction in post hoc mistranslation rates is particularly evident on the toxin-antitoxin (E3) landscape at the smallest population size (figure S9a, blue line).

On the toxin-antitoxin (E2) landscape, we observe an increase in post hoc mistranslation rates in walks without mistranslation after the first $10^3$ mutations (figure S9b, blue line). This increase happens because the outcome of adaptive walks on the toxin-antitoxin (E2) landscape is biased towards genotypes with higher mistranslation rates: At a population size of $N = 10^4$, 65% of walks end at two amino acid sequences with slightly elevated post hoc mistranslation rates (1.524% and 1.525% mistranslated). In contrast, on the toxin-antitoxin (E3) and antibody-binding landscapes, the two most common amino acid sequences at the end of the adaptive walks account for 30% and 9% of walks respectively. Consequently, unless there are strong outliers, as is the case for the toxin-antitoxin (E2) landscape, a long-term reduction in post hoc mistranslation rates occurs.

#### 8 Approximating mistranslation-free fitness

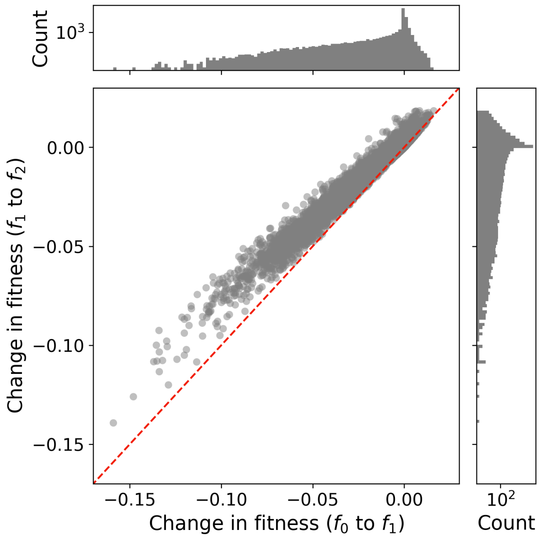

Figure S10: Changes in fitness due to mistranslation correlate strongly between inferred mistranslation-free fitness values ($f_0$) and the predictions of our model ($f_2$). Main panel: Difference in fitness between inferred fitness values ($f_0$) and empirical fitness measurements ($f_1$) of all polypeptide sequences in the antibody-binding landscape (horizontal axis), and difference in fitness between the empirical fitness measurements ($f_1$) and the fitness predicted by our model ($f_2$). The model predictions ($f_2$) are an average of the fitness of all synonymous nucleotide sequences encoding a given polypeptide sequence. The diagonal red dashed line denotes equivalent changes in fitness between $f_0$ to $f_1$, and $f_1$ to $f_2$. Upper panel: Distribution of the differences in fitness between inferred fitness values $f_0$ and empirical fitness measurements $f_1$. Right panel: Distribution of the differences in fitness between empirical fitness measurements $f_1$ and model predictions $f_2$.

For most of this contribution, we assume that the empirical fitness measurements $f_1$ are free of the effect of mistranslation and simulated the effect of mistranslation on these measurements to calculate fitness with mistranslation $f_2$. We will now estimate the true fitness of these genotypes $f_0$ from the empirical measurements $f_1$. Because the available data report fitness $f_1$ at the level of amino acids and not nucleotide sequences, we can only approximate the effect of mistranslation on fitness measurements when trying to infer $f_0$.

We estimate mistranslation rates at the level of amino acid sequences, using experimentally measured mistranslation rates for each codon that encodes a specific amino acid sequence [4], and averaging the mistranslation rate over all such codons. With these averaged mistranslation rates we estimate the probability that translation produces either the encoded amino acid sequence or an alternative sequence. For every amino acid sequence in the antibody-binding landscape, we calculate the probability of translation producing the encoded amino acid sequence, or all other amino acid sequences in the landscape. We place these probabilities into a matrix $A$ of size $n \times n$, where $n = 149361$ is the number of amino acid sequences in the antibody-binding landscape. We then infer $f_0$ by solving the system of linear equations $f_0 = Af_1$.

The inferred fitness $f_0$ has a mean of $8.15 \times 10^{-2} \pm 0.402$, which is significantly different to the mean fitness of $f_1$ $8.05 \times 10^{-2} \pm 0.395$ and the mean fitness of $f_2$ $8.05 \times 10^{-2} \pm 0.391$ (Kruskal-Wallis $H = 2530$, $p \ll 10^{-300}$, Dunn's test with Bonferroni correction, all pairwise comparisons are significant with $p \leq 1.06 \times 10^{-70}$). The predicted changes in fitness from $f_0$ to $f_1$ are qualitatively similar to the predicted changes from $f_1$ to $f_2$. Indeed, the changes in fitness between $f_0$ and $f_1$ are correlated with the changes in fitness between $f_1$ and $f_2$ (Kendall's $\tau = 0.477$, $p \ll 10^{-300}$, $n = 149361$, figure S10).
